# Loss of killifish cGAS/STING function attenuates cellular senescence and age-related signatures but does not extend organismal life span

**DOI:** 10.1101/2024.10.08.617203

**Authors:** Eugen Ballhysa, Roberto Ripa, Nadine Hochhard, Tin Tin Manh Nguyen, Jennifer Brazell, Baptiste Ferreri, Elena Hoffmann, Joachim Steiner, Adam Antebi

## Abstract

The cGAS/STING pathway is a central innate immune signaling pathway whose chronic activation has been implicated in numerous age-related pathologies, yet its impact on life span itself is unknown. Here we engineered knockouts of this pathway in the killifish *Nothobranchius furzeri*, and assessed physiology and aging. *In vitro*, loss of killifish cGAS or STING mitigated DNA damage-induced senescence in cultured fibroblasts. *In vivo*, cGAS knockout unexpectedly led to low-grade inflammation. It also attenuated changes in gene expression in response to DNA damage in young animals, and age-related changes in the old, suggesting dampening of senescence and aging. Necroscopy indicated that tissue pathology appeared milder overall in both mutants, though some tissues showed enhanced sterile macrophage infiltration. Despite an attenuated aging signature, however, longevity was not significantly different from wild type. Our findings reveal a potential tradeoff, where inhibiting the cGAS/STING pathway alleviates age-related signatures, but increases sterile inflammation, offsetting beneficial effects on lifespan.

## Introduction

Aging is often accompanied by low grade chronic inflammation, termed inflammaging [1], which contributes to age-related pathology and decline of tissue homeostasis. Many age-related diseases, including cardiovascular [2], neurodegenerative [3, 4], cancer [5] and metabolic syndromes [6] are associated with increased, unresolved inflammation, which can interfere with regenerative processes necessary for cellular repair and exacerbate disease. Hence, tempering chronic inflammation is one avenue to improve health into old age.

A key source of inflammaging is the nucleic acid sensing innate immune signaling pathway cGAS/STING [7]. This pathway detects DNA in the cytosol [8] – arising from DNA damage [9–11], mitochondrial dysfunction [12, 13], or infection [14] – which stimulates the activity of the cyclic GMP-AMP-synthesizing enzyme cGAS to produce the second messenger, 2’3’- cGAMP (cGAMP) [15, 16]. Subsequently, cGAMP binds to its sensor STING, which activates regulatory cascades to promote expression of type I interferon response genes and other inflammatory cytokines [17]. While this pathway is important for innate immune protection, its chronic stimulation can lead to age related pathology [18].

The cGAS-STING pathway also plays a central role in cellular senescence, whereby cells inflicted with DNA damage [11, 19], mitochondria dysfunction [12, 20], and other cellular harm [9], permanently arrest in the cell cycle and secrete inflammatory cytokines, chemokines and matrix remodeling proteins, collectively termed the senescence associated secretory phenotype (SASP) [21]. Cytosolic DNA present in senescent cells is typically detected by the cGAS/STING pathway, which then triggers the SASP. Continuous unresolved damage can act in an endocrine/paracrine fashion to foster chronic systemic inflammation [22]. Hence, limiting the activity of cGAS-STING pathway during aging seems like a viable approach to mitigate chronic inflammation and improve health outcomes. Indeed, late life administration of STING inhibitors can ameliorate neurodegeneration and slow cognitive decline in mice [3].

Despite its importance in age-related disease, the impact of cGAS/STING pathway on life span itself remains unstudied. Here we chose to elucidate the role of the cGAS/STING pathway in the short-lived African turquoise killifish *Nothobranchius furzeri*, an emerging model system for studying aging. The *N. furzeri* GRZ strain lives a mere 4-8 months, yet exhibits many features of mammalian aging and age-related pathology [23]. In this work we used CRISPR-engineering to knockout killifish cGAS and STING and study their function in senescence and aging. We hypothesized that inhibition of this pathway could lead to significant health benefits manifest as an increase in longevity. As with mammals, we found that diminishing this pathway mitigates expression of cellular senescence markers induced by DNA damage *in vitro* and *in vivo*. Further, it attenuates age-related transcriptional changes, and tends to reduce age-related pathology. Unexpectedly, however, knockout mutants showed elevated inflammation and a similar life span as wild type, suggesting there may be tradeoffs to any benefits and that this pathway is not limiting for organismal vitality.

## Results

### cGAS structure and function *in vitro* are conserved from teleosts to mammals

The cGAS/STING pathway has been extensively studied in mammalian cell culture models, but relatively little is known about its function in teleosts [24–27]. We therefore sought to investigate its function in the short-lived African killifish *N. furzeri*, an important model system for vertebrate aging [28–30]. BLAST analysis of the *N. furzeri* genome (UI_Nfuz_MZM_1.0 reference genome) revealed one copy of the cGAS gene (XM_015944714.2, kcGAS), showing 35% amino acid identity to its human ortholog (Figure 1a, b). Like other members of the family, the predicted protein contained conserved domains including the NTase core, involved in the catalytic production of cGAMP, the mab21 domain, implicated in DNA binding, as well as a non-conserved N’-terminal region (Figure 1a). Structural alignment of the human cGAS (PDB 5VDO) with AlphaFold 3-predicted model of kcGAS showed a root mean square deviation of 1.015 angstroms across 277 pruned atom pairs (Extended Data Figure 1a), indicating a high degree of tertiary structural similarity.

**Figure 1.**
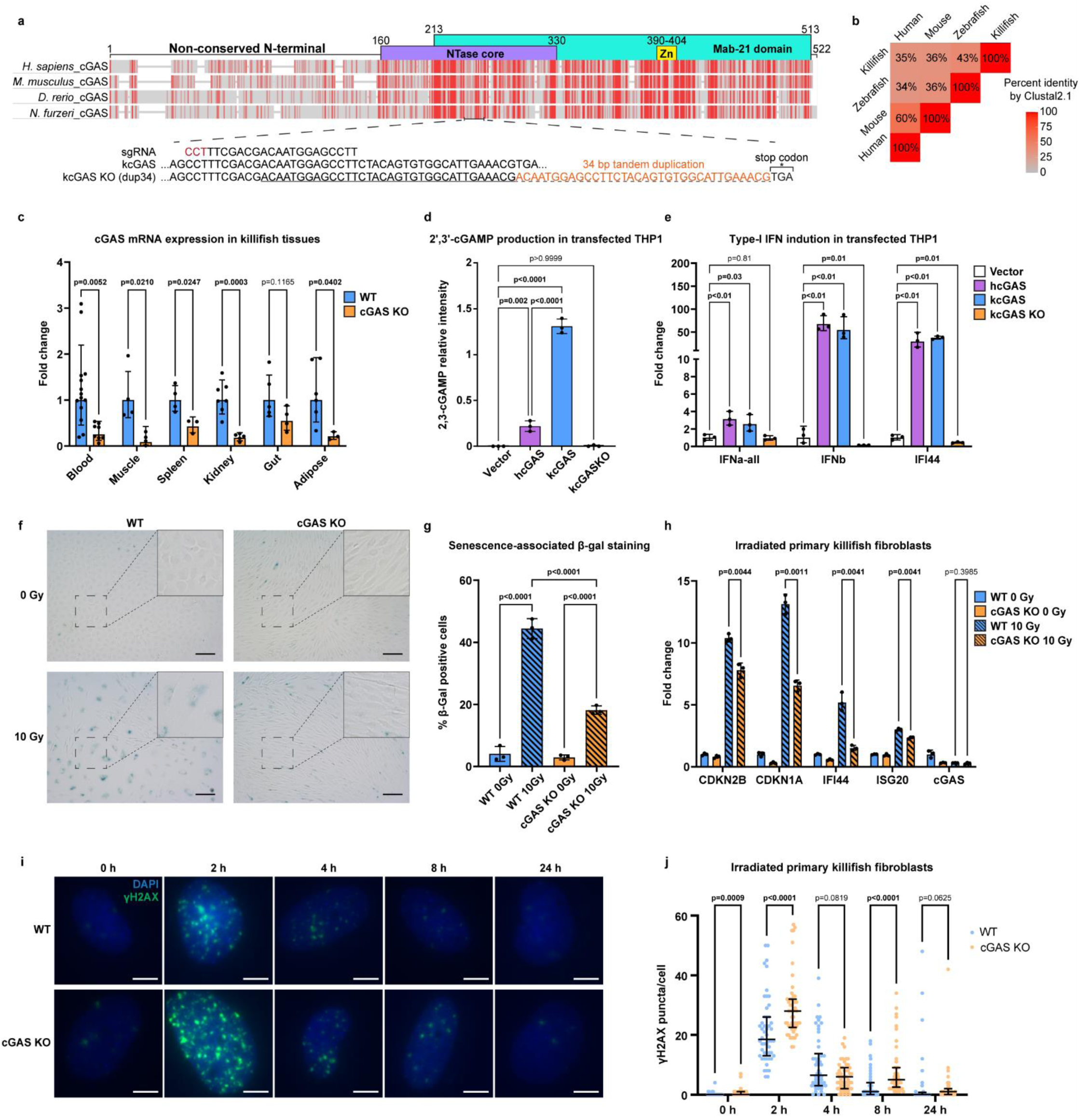
cGAS structure and function are conserved from teleosts to humans. **a**, Multiple sequence amino acid alignment of cGAS with orthologues from other species using constraint-based multiple sequence alignment tool (COBALT). Red= identical amino acids, grey= non-identical. Above the alignment are shown the NTAse and Mab21 domains and the Zn finger. Below the alignment is a schematic of the CRISPR-Cas9-generated mutated region of killifish cGAS and the used sgRNA. **b**, Triangle heatmap showing percent identity between aligned cGAS genes of different organisms using ClustalW. **c**, Expression of cGAS mRNA measured using qPCR in different tissues of WT and cGAS KO mutant killifish. Each dot represents data from one fish. n ≥ 3 per tissue and genotype. Statistical analysis was done with Student’s t-test using the normalized expression values of cGAS to GAPDH. **d**, LC–MS-based targeted metabolomics of 2’,3’-cGAMP extracted from THP1 cells transfected with human cGAS (hcGAS), killifish cGAS (kcGAS), and mutated killifish cGAS (kcGAS KO). Statistical analysis was performed using one-way ANOVA with Dunn–Šidák correction for multiple comparisons. **e**, Expression of interferon and interferon-stimulated genes by qPCR from cell extracts of cGAS KO THP1 cells transfected with the indicated cGAS genes. Samples are identical to those used in (D). Statistical analysis was done using Student’s t-test using the normalized expression values to GAPDH. **f**, Bright field images of WT and cGAS KO killifish primary fibroblasts 9 days post 10 Gy γ-radiation, stained for senescence associated β-galactosidase activity. Scale bar = 100 μm. **g**, Quantitation of senescence associated β-galactosidase positive cells from images in (**f**). At least 100 cells were counted per replicate. Statistical analysis was performed using One-way ANOVA with Dunn–Šidák correction for multiple comparisons. **h**, qPCR measurement of senescence and interferon-stimulated genes in irradiated WT and cGAS KO primary fibroblasts. Statistical analysis was done with Student’s t-test using the normalized expression values to EIF3c. **i**, Representative images of primary killifish fibroblasts stained for γH2AX (Green) and DAPI (blue) of indicated genotypes. Cells were fixed at the indicated times after 10 Gy of γ-radiation. Non-irradiated cells were used as 0 h controls. Scale bar = 20 μm. **j**, Quantitation of phosphorylated γH2AX puncta measured from images in (**i**). Statistical analysis was performed using the Mann-Whitney test. Each dot represents one cell, n ≥ 50 cells were measured for each condition. All cell culture experiments in **d**-**j** were performed in 3 independent biological replicates with n = 3 plates each time. Each panel shows one of three independent experiments.

To begin to unravel the physiologic function of kcGAS, we first generated a CRISPR/Cas9 knock-out. The sgRNA caused random mutagenesis within the NTase core, resulting in a 34bp tandem duplication that led to a frameshift and early stop codon, and hence a presumptive null allele (Figure 1a) [31]. Analysis of kcGAS mRNA expression levels revealed strong downregulation in multiple tissues, indicating nonsense-mediated RNA decay (Figure 1c). Furthermore, by LC-MS we previously observed a complete lack of cGAMP in the liver and gut of the cGAS KO fish [31]. All together these results confirm the efficacy of the knock-out.

Upon activation, cGAS triggers the initiation of type-I interferon signaling in mammalian systems [17]. To validate the conservation of the axis in the killifish we initially adopted a heterologous cell culture setting. We transfected wild-type killifish cGAS (kcGAS), mutated killifish cGAS (kcGAS KO) and human cGAS (hcGAS) into cGAS deficient human THP1 cells using an expression vector carrying GFP as an internal control for transfection efficiency (Extended Data Figure 1b). We detected high levels of cGAMP in cells expressing human and killifish cGAS, while the empty vector and kcGAS KO showed no cGAMP production (Figure 1d). Interestingly, kcGAS produced higher levels of cGAMP compared to hcGAS (Figure 1d), possibly reflecting higher intrinsic enzymatic activity of kcGAS, similar to mouse cGAS [32]. Indeed, despite higher levels of cGAMP, kcGAS had lower transfection efficiency compared to hcGAS (Extended Data Figure 1c, d) while transgene expression load was similar within transfected cells (Extended Data Figure 1e, f). We also observed that transfection efficiency was lower in WT kcGAS and hcGAS compared to kcGAS KO (Extended Data Figure 1c-f), presumably because of activation of the cytosolic DNA response. In line with this, the presence of kcGAS but not kcGAS KO was sufficient to stimulate expression of type I interferon response genes (Figure 1e). We conclude that kcGAS contains intrinsic cGAMP producing activity that is disrupted by the kcGAS KO mutation.

Concurrently, we also investigated the role of killifish STING (kSTING). The killifish genome harbors one functional copy of kSTING, which showed 36% percent identity to human STING1 (hSTING) (Extended Data Figure 2a, b). Alignment of the AlphaFold 3-predicted kSTING model with hSTING dimer (PDB 8FLM) showed a root mean square deviation of 1.28 angstroms from 164 pruned atom pairs (Extended Data Figure 2c), suggesting a high degree of conservation in tertiary structure. We next generated a killifish STING KO mutant strain using CRISPR/Cas9 (Extended Data Figure 2a), in this case using two sgRNAs simultaneously, and obtained two small deletions of 2 and 5 base pairs, both of which led to frameshift mutations. qPCR confirmed a marked reduction in STING mRNA levels in tissues of this mutant, indicating non-sense mediated decay (Extended Data Figure 2d). Transfection of kSTING gene into STING KO THP1 monocytes, unlike kcGAS, failed to activate immune genes (Extended Data Figure 2e), conceivably due to incompatibility with heterologous human co-factors such as TBK1 (Extended Data Figure 2F) [33–35]. In addition, transfection efficiency in STING KO THP1 cells was markedly lower (Extended Data Figure 2g, h) compared to cGAS KO cells (Extended Data Figure 1c-f), possibly because of the presence of endogenous cGAS.

In mammalian cell culture, cGAS and STING are required to establish aspects of the senescent phenotype [3, 9, 11, 19]. Whether this function is conserved in teleosts, however, remains unknown. We therefore asked if killifish cGAS plays a similar role in cellular senescence. To test this idea, we isolated primary fibroblasts from the fins of WT and cGAS KO killifish and induced senescence with DNA damage, subjecting cell cultures to 0 Gy or 10 Gy of γ-radiation. Nine days post-irradiation, we observed that cGAS KO fibroblasts retained their fiber-like morphology and had significantly fewer cells staining for senescence associated β-galactosidase activity compared to WT fibroblasts (Figure 1f, g), suggesting lower levels of senescence. From these cell cultures, we extracted RNA and performed qPCR to measure the expression of senescence markers (CDKN2B, CDKN1A) and IFN signaling genes (IFI44, ISG20). While all genes showed induction post-irradiation, cGAS KO fibroblasts had significantly lower expression levels compared to WT fibroblasts (Figure 1h). We also established killifish STING KO primary fibroblasts and carried out the same experiment. Nine days post-irradiation, STING KO fibroblasts also exhibited blunted expression of senescence and interferon markers (Extended Data Figure 2i). Notably, this experiment complemented the heterologous cell culture data described above and showed the ability of kSTING to fully promote the interferon response in its native cellular environment. Altogether, these data support the evidence that killifish cGAS and STING loss of function lead to mitigation of cellular senescence in an evolutionary conserved manner.

To further understand the response to the genotoxic stress, we irradiated WT and killifish cGAS KO primary fibroblasts and stained them for γH2AX at various time points (0h, 2h, 4h, 8h, and 24h). Even without irradiation (0h) cGAS KO fibroblasts showed few but significantly more γH2AX puncta compared to WT cells. At 2h post-irradiation, γH2AX puncta spiked higher in cGAS KO cells compared to WT cells, and persisted over the next few hours until they dropped to basal levels 24h post-irradiation (Figure 1i, j). This finding suggests that, despite dampened senescence and immune marker activation, cGAS KO primary fibroblasts inherently display more DNA damage and/or take longer to repair such damage.

### Killifish cGAS impacts innate immunity regulation *in vivo*

We next wondered how lack of cGAS affected killifish physiology *in vivo* under basal conditions. To investigate this, we first performed bulk RNAseq from the kidneys of young (7-8 week) healthy WT and cGAS KO fish. We first chose the kidney because of its central role in hematopoiesis, immune cell production and differentiation, and its function as the primary lymph node in teleosts [36].

Principle component analysis (PCA) plots showed subtle differentiation of the genotypes (Figure 2a) and minus-average (MA) plot revealed only modest overall transcriptomic differences (Figure 2b). Among the changed transcripts were several involved in vesicular trafficking (ARF1L, SREBF1, SEC23b, SEC24c), secreted metalloproteinases (MMP19, MEP1a.1), TSC1b (mTOR signaling inhibitor) and several lncRNAs. To decipher potential differences further, we used gene set enrichment analysis (GSEA) and identified a number of significantly enriched pathways in the absence of cGAS. Notably, immune-related pathways such as Rig-I-like receptor signaling, Toll-like receptor signaling, Nod-like receptor signaling, and complement and coagulation cascades, were upregulated in the cGAS KO relative to wild type (Figure 2c). Closer analysis of these upregulated immune genes revealed that some act directly downstream of cGAS, such as STING and TBK1, suggesting compensation for cGAS loss (Figure 2d). However, the majority of these upregulated immune genes act upstream (TRAF6, RIPK1L), downstream (IL12BB) or within NF-kB signaling cascades (NFKB1, RELA, TNFA), even in the absence of an overt stimulus (Figure 2d) suggesting increased inflammation. Other gene categories upregulated in cGAS KO included lysosome function and apoptosis. Upregulation of lysosomal genes (CTSZ, CTSB, ATP6V0C and ATP6V0B) suggests an enhanced capacity for degradation and recycling of cellular components, which is a common response to cellular stress or damage, while upregulation of apoptotic genes (CASP7, MAP3K14, BAX and BIRC2) suggests an increase in apoptotic signaling, which might indicate that mutant cells are under stress and are more prone to undergo cell death.

**Figure 2.**
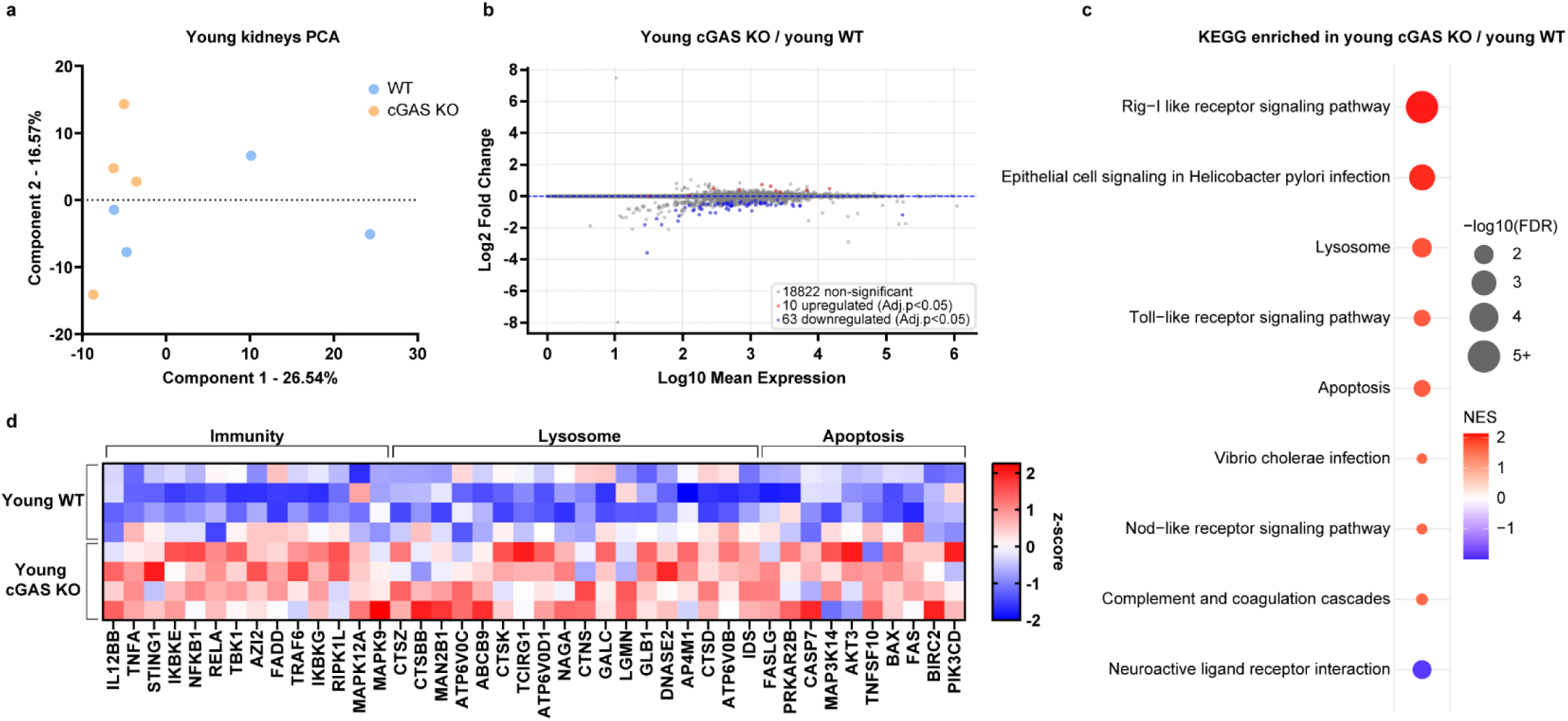
Loss of cGAS stimulates upregulation of downstream innate immune signaling components in young killifish. **a**, PCA plot of WT and cGAS KO transcriptomes from young (8 weeks) killifish kidneys. b, MA plot showing differential mRNA expression comparing cGAS KO to WT young kidneys. Throughout the paper, all genes with adjusted p-value < 0.05 are considered significant and marked as upregulated (red) or downregulated (blue). **c**, GSEA comparing transcriptomes of young kidneys of cGAS KO to WT. All significantly upregulated and downregulated KEGG pathways are shown (FDR < 0.05). Unless noted otherwise, gene sets throughout the paper are derived from the KEGG_LEGACY subset of canonical pathways. Additional KEGG pathways and associated genes throughout are found in our data repository (see Additional information). **d**, Heatmap of z-score normalized expression values of the genes with the highest positive and lowest negative rank metric scores from representative GSEA pathways shown in (**c**). NES: Normalized enrichment score, FDR: False discovery rate. N = 4 fish per genotype.

To examine another tissue, we also performed RNAseq of gut samples from young WT and cGAS KO killifish. We chose the gut since this tissue serves as a first line of defense against various pathogens, in line with the innate immune function of cGAS/STING. As with the kidneys, we saw no clear separation of genotypes in the PCA plot, nor was there a large impact on gene regulation (Extended Data Figure 3a, b). GSEA revealed processes that were significantly upregulated (complement, ribosome) and downregulated (DNA replication, cell cycle, amino acyl tRNA biosynthesis) in cGAS KO guts relative to WT (Extended Data Figure 3c), although we observed high variability between replicates in core regulatory genes within these processes (Extended Data Figure 3d).

We also carried out transcriptome analysis comparing young WT and STING KO kidneys under basal conditions. This analysis revealed only minor differences in overall transcription and little impact on cellular pathways (Extended Data Figure 3e-g).

### Killifish cGAS modulates the transcriptional response to DNA damage and senescence *in vivo*

Given the small differences we observed in young animals under basal conditions, we next asked how the cGAS/STING pathway responded to stress *in vivo*. In particular, since we had observed that cGAS impacts DNA damage response and senescence *in vitro*, we wished to examine these features *in vivo*. To this end, we irradiated young healthy WT and cGAS KO fish with 15Gy of γ-radiation and 5 days later, harvested their tissues and assessed the transcriptional response in the kidneys. We chose this time point in order to potentially capture early senescence events *in vivo* [37, 38]. In this case, PCA plots showed a clear separation between irradiated (15 Gy) and non-irradiated (0 Gy) samples (Figure 3a) with hundreds of genes responding to the stimuli (Figure 3b) and widespread transcriptional regulation dependent on cGAS (Figure 3a, c). This indicates that cGAS is important for transcriptional regulation in response to DNA damage in young healthy tissues.

**Figure 3.**
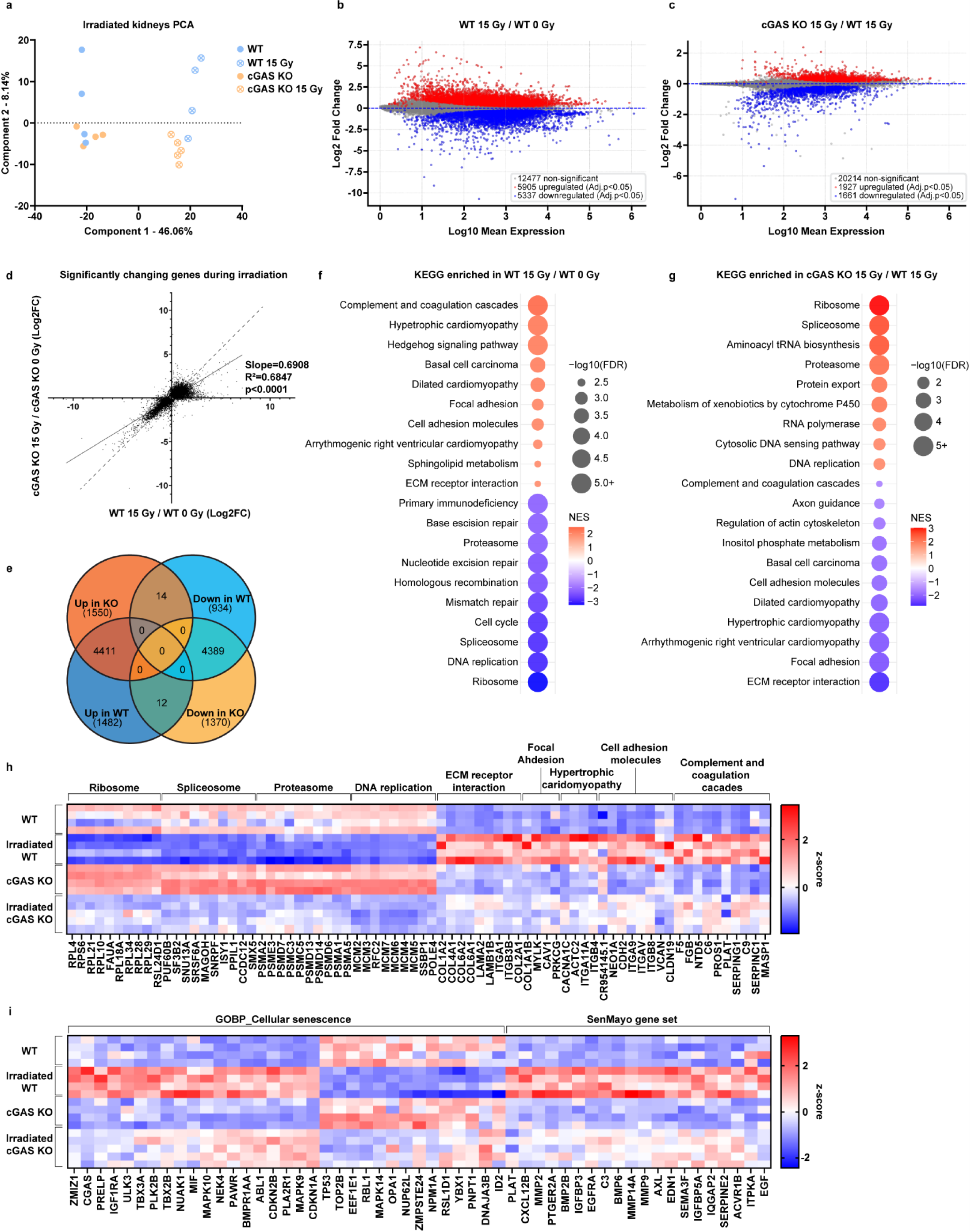
cGAS modulates DNA damage induced transcriptional changes in vivo. **a**, PCA plot of WT and cGAS KO transcriptomes from kidneys of young (8 weeks) killifish, which were either irradiated or non-irradiated. Kidneys were harvested 5 days after irradiation. **b**, MA plot showing differential mRNA expression comparing 15 Gy irradiated to non-irradiated young WT kidneys. All genes with adjusted p-value < 0.05 are considered significant, upregulated (red), downregulated (blue). **c**, MA plot showing differential mRNA expression comparing irradiated cGAS KO to irradiated WT young kidneys. All genes with adjusted p-value < 0.05 are considered significant, upregulated (red), downregulated (blue). **d**, Linear regression analysis of the Log2(fold change) in all genes significantly changed in expression during irradiation in WT and cGAS KO kidneys, showing a slope significantly < 1. A dashed line with slope = 1 is shown for comparison. **e**, Venn diagram depicting the overlap of significantly changed genes post-irradiation in WT and cGAS KO kidneys. **f**, GSEA comparing irradiated to non-irradiated WT kidneys. The top KEGG pathways upregulated and downregulated in irradiated WT are shown. **g**, GSEA comparing irradiated cGAS KO and WT kidneys. The top KEGG pathways upregulated and downregulated in cGAS KO are shown. **h**, Heatmap of z-score normalized expression values of the genes with the highest positive and lowest negative rank metric scores from representative GSEA pathways shown in (**f**, **g**). **i**, Heatmap of z-score normalized expression values of genes with the highest positive and lowest negative rank metric scores from GSEA when comparing irradiated to non-irradiated WT kidneys. The gene sets investigated were the SenMayo gene set and the GOBP Cellular Senescence gene set. GOBP: Gene Ontology Biological Process. N >= 4 fish per genotype and treatment

Interestingly, linear regression of all significantly regulated genes of WT and cGAS KO during irradiation showed high linear correlation (R^2^ = 0.6847) but with a slope significantly less than 1 (m = 0.6908). This observation indicates that genes are generally regulated in the same direction, but with a lower response in cGAS KO kidneys during irradiation compared to WT (Figure 3d). Comparing transcripts up- and downregulated in WT and cGAS KO showed high overlap as well as distinct significantly changing gene sets in WT and cGAS KO kidneys, identifying gene responses completely dependent on cGAS (Figure 3e). Virtually no genes were oppositely regulated, further supporting the idea that cGAS modulates the magnitude rather than the directionality of the transcriptional response.

To better understand which pathways were differentially regulated in the absence of cGAS, we performed GSEA. In line with the linear regression analysis, the regulation of most pathways by irradiation was significantly dampened in the cGAS KO kidney transcriptome (Figure 3f, g). These included pathways involved in innate immunity (complement components C6, C9, SERPING1), extracellular matrix, cell adhesion and focal adhesion (COL1A2, LAMA2; CDH, ITGA9; CAV1, MYLK) and cardiomyopathy (ITGB4, ACTC2) (Figure 3g, h). In addition, we observed that the cGAS KO kidney transcriptome showed higher basal levels of processes associated with growth and proliferation, which were less downregulated by irradiation (Figure 3h). With this in mind, we revisited our data on young non-irradiated cGAS KO versus WT kidneys, and noticed that a number of specific genes associated with DNA replication (e.g. MCM family), protein homeostasis (e.g. proteasome subunits PSMA2, PSME3; ribosomal subunits RPS6, RPL21) and mRNA splicing (e.g. PUF60B, SF3B2) were elevated relative to WT (Figure 3h), though their pathways did not emerge as significantly enriched (Figure 2c). Conceivably, this upregulation may reflect an attenuated stress response or relaxation of checkpoints that regulate growth and cell division.

Curiously, the expression of DNA repair pathways themselves were downregulated in both cGAS KO and WT at this 5-day timepoint consistent with previous observations that cells on a senescent trajectory downregulate DNA repair [39]. Because the KEGG_LEGACY pathways only partially represent senescence and SASP factors, we decided to specifically investigate the expression of genes under the Cellular Senescence GO Biological Process (GOBP) term as well as the SenMayo [40] gene set. Notably, we observed a blunted senescent gene expression in the cGAS KO kidneys (Figure 3i) especially of SASP components (MIF, CXCL12B, IGFBP3/5A, MMP2/9), corroborating our *in vitro* findings.

Altogether, despite the presence of a DNA damage and senescence responses in the absence of cGAS, the magnitude of regulation was markedly reduced, suggesting that cGAS serves to amplify the transcriptional response of these processes *in vivo*.

We also analyzed the gut transcriptome from the same irradiated fish as above. As with the kidney, PCA plots showed that the genotypes clustered separately after irradiation (Extended Data Figure 4a). Linear regression of the significantly changed genes during irradiation comparing both genotypes again showed a decreased slope (m=0.4059), though the linear correlation was weaker (R^2^=0.3564) (Extended Data Figure 4b). GSEA analysis revealed widespread regulation of multiple pathways after irradiation (Extended Data Figure 4c), while the lack of cGAS again blunted the response, especially of ribosomal protein regulation (Extended Data Figure 4c, d). Notably, the attenuation of gene and KEGG pathway regulations were less prominent than those observed in the kidney. However, senescence and SASP activation were blunted similarly to the kidney (Extended Data Figure 4e), suggesting cGAS functions in multiple tissues during genotoxic stress, but the breadth of regulation appears tissue specific.

We next investigated the transcriptional response to DNA damage in the STING KO kidneys. PCA plots showed that irradiated STING KO transcriptomes separated from irradiated WT, and a number of genes were differentially regulated (Extended Data Figure 4f, g). In this case, the attenuation of gene regulation, as seen in the cGAS KO, was not observed in the STING KO (Extended Data Figure 4h). In fact, pathways that were most affected, namely the complement and coagulation cascades and proteasome (Figure 3f), were more upregulated or downregulated, respectively (Extended Data Figure 4h, i). By comparison, proteasome subunits did not show uniform regulation across batches when relating non-irradiated to irradiated WT kidneys (Figure 3h, Extended Data Figure 4i). Still, genes involved in inflammation (C7, C4B and SERPINA1) and cellular stress responses (SCARA3 and NTD5) showed upregulation in the STING KO, opposite to what we observed with cGAS KO (Figure 3h). While many senescent genes appeared to be similarly regulated between WT and STING KO, we noticed a number of SASP genes within the SenMayo gene set (IGFBP family, CXCL12B) to have blunted regulation (Extended Data Figure 4i), which is consistent with what has been observed in human cell cultures [3, 9, 19].

Collectively, these observations hint towards a broad STING-independent and tissue specific role of cGAS as an important modulator of the transcriptional response to DNA damage and senescence.

### Killifish cGAS modulates the aging transcriptome

Aging is associated with increased DNA damage, inflammation, and senescence. In addition, aged mice brains have shown higher cGAMP levels when compared to young brains, indicating higher cGAS activity or lower cGAMP turnover with age [3]. In killifish we detected increased cGAMP levels in the old kidneys, but no significant change in the gut when compared to young tissues (Extended Data Figure 5a). Hence, we sought to understand the impact of cGAS to the normative aging transcriptome. We used the young (8 weeks) healthy WT and cGAS KO fish as reference and performed RNAseq in old (18 weeks) WT and cGAS KO kidneys. Similar to what we observed post-irradiation, PCA plots showed that the two genotypes separated better in old age than in young (Figure 4a). Overall, the killifish kidney transcriptome was vastly deregulated, and the aged cGAS KO transcriptional profile deviated significantly from the aged WT counterpart (Figure 4b, c).

**Figure 4.**
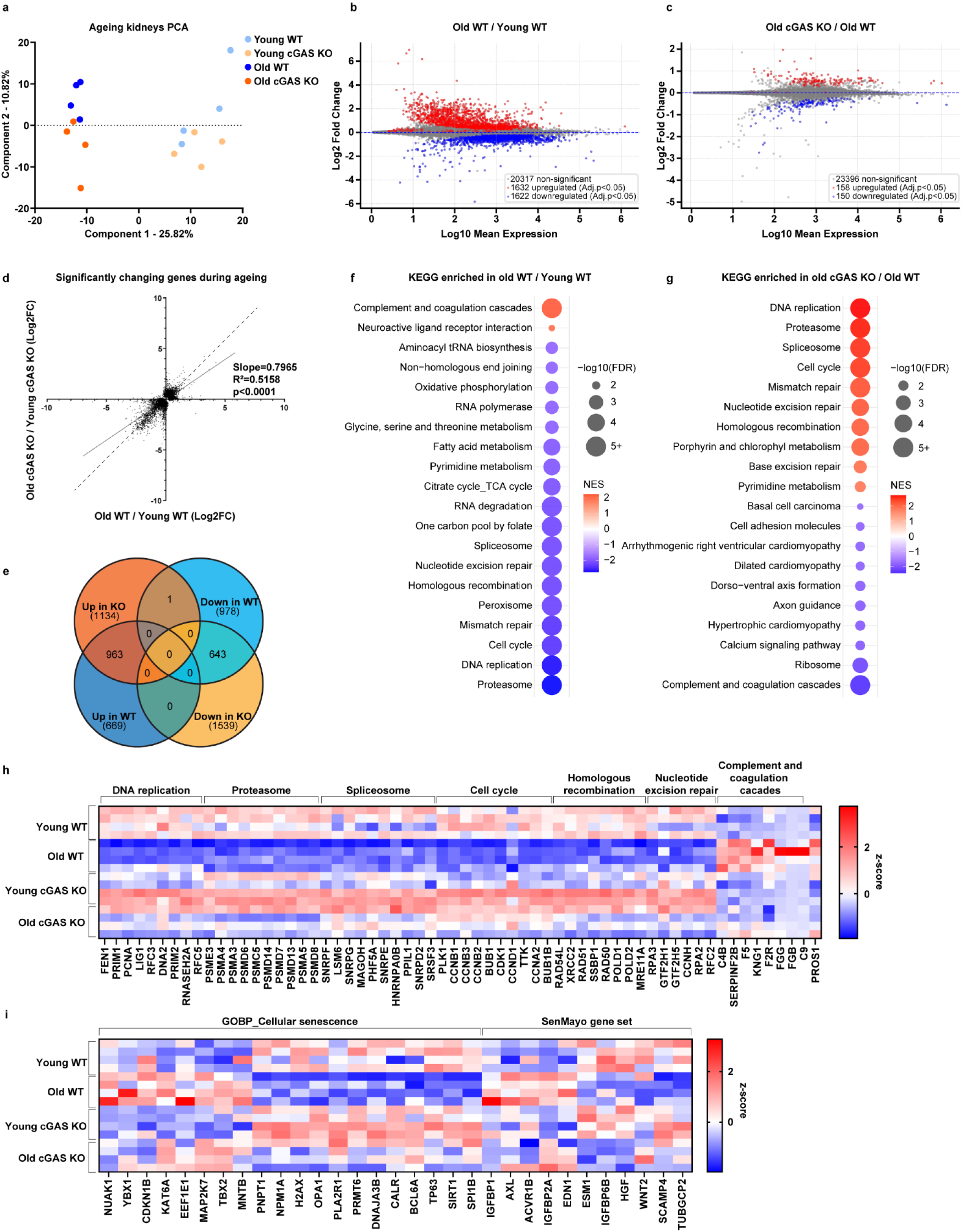
cGAS modulates transcriptional changes during ageing. **a**, PCA plot of WT and cGAS KO transcriptomes from young (8 weeks) and old (18 weeks) killifish kidneys. **b**, MA plot showing differential mRNA expression comparing old to young WT kidneys. All genes with adjusted p-value < 0.05 are considered significant, upregulated (red), downregulated (blue). **c**, MA plot showing differential mRNA expression comparing old cGAS KO to old WT kidneys. All genes with adjusted p-value < 0.05 are considered significant, upregulated (red), downregulated (blue). **d**, Linear regression analysis of the Log2(Fold change) of all genes significantly changed in expression during ageing in WT and cGAS KO kidneys, showing a slope significantly < 1. Genes significantly regulated only in WT ageing or cGAS KO ageing or inversely regulated are also included. A dashed line with slope = 1 is shown for comparison. **e**, Venn diagram depicting the overlap of significantly changed genes during ageing in WT and cGAS KO kidneys. **f**, GSEA comparing old to young WT kidneys. All KEGG pathways significantly upregulated and the top downregulated in old WT kidneys are shown. **g**, GSEA comparing old cGAS KO to old WT kidneys. The top KEGG pathways upregulated and downregulated in cGAS KO are shown. **h**, Heatmap of z-score normalized expression values of the genes with the highest positive and lowest negative rank metric scores from representative GSEA pathways shown in (**f**, **g**). **i**, Heatmap of z-score normalized expression values of genes with the highest positive and lowest negative rank metric scores from GSEA when comparing old to young WT kidneys. The gene sets investigated were the SenMayo gene set and the GOBP_Cellular Senescence gene set.

As with irradiation, cGAS clearly impacted the aging kidney transcriptome. Comparing cGAS KO to WT, linear regression of all transcripts significantly regulated with age showed a significantly reduced slope (m=0.7965) (Figure 4d), with reasonable linear correlation (R^2^=0.5158), and virtually no genes inversely regulated in the two genotypes (Figure 4e). These findings point towards robust changes in gene regulation with age, which are blunted by cGAS KO. GSEA revealed that KEGG pathways upregulated with normative aging included complement and coagulation cascades, and neuroactive ligand receptor interaction, while KEGG pathways downregulated with aging included proteasome, DNA replication, cell cycle, DNA repair, peroxisome, spliceosome, one-carbon metabolism, TCA cycle and oxphos among others (Figure 4f).

Attenuation of gene expression changes by cGAS KO applied broadly to a majority of pathways dysregulated with age, revealing a widespread effect (Figure 5f, g). In particular, GSEA revealed that DNA replication and repair pathways, cell cycle, spliceosome, proteasome, and pyrimidine metabolism were less downregulated (Figure 4f, g), while complement and coagulation and ribosome were less upregulated in old cGAS relative to old WT. For example, the aging cGAS KO kidneys lacked the strong reduction of cell proliferation markers (FEN1, PCNA, RFC3, CCNB1/2/3), suggesting less replicative arrest (Figure 4h). These patterns suggest that cGAS KO cells may be more proliferative and actively engaged in the cell cycle.

**Figure 5.**
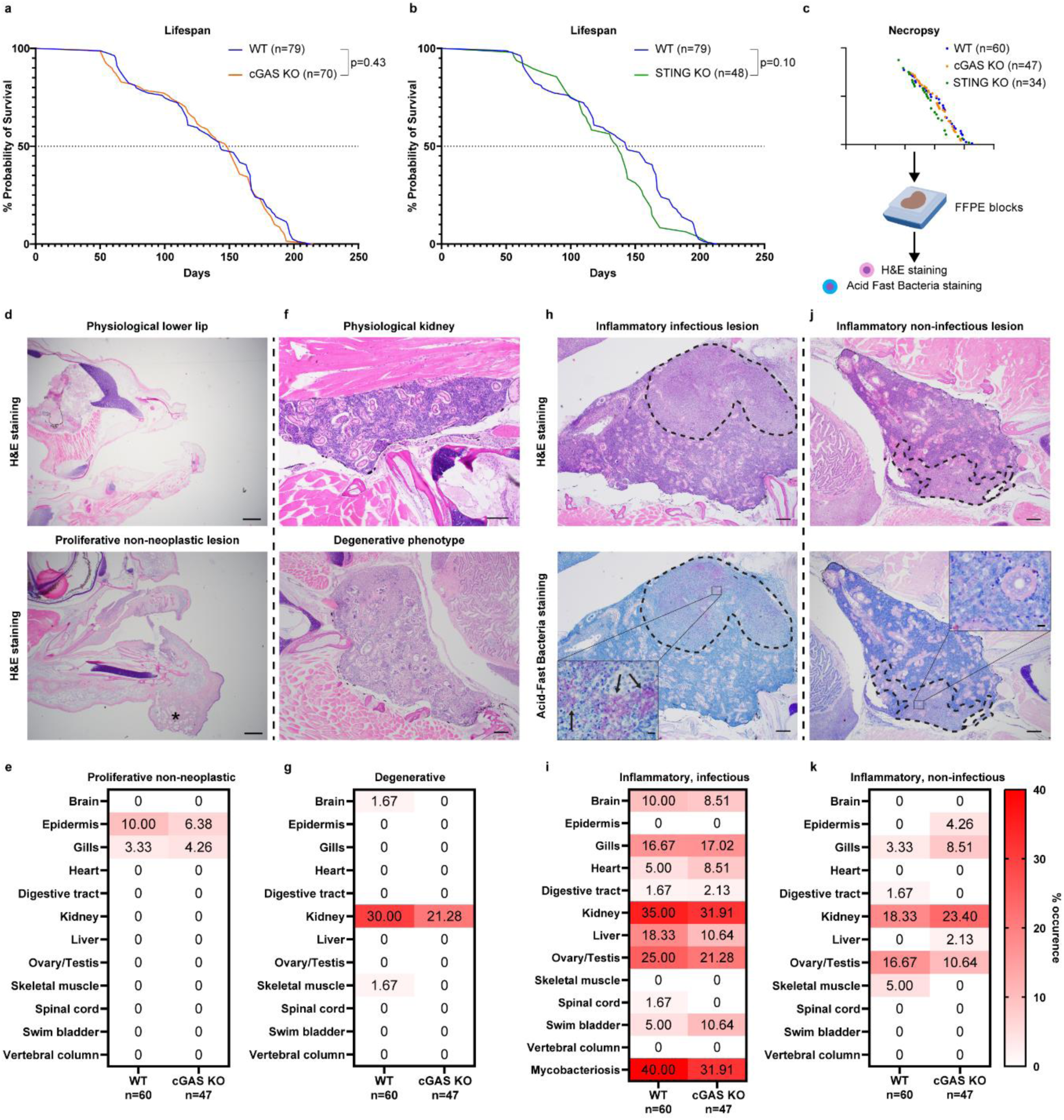
Killifish cGAS/STING pathway influences degenerative and inflammatory disease occurrence but not lifespan. **a**. Kaplan-Meier curve showing the survival of WT (n = 79) and cGAS KO (n = 70) killifish. Indicated p-value represents comparison between cGAS KO and WT using the log-rank Mantel-Cox test. **b**, Kaplan-Meier curve showing the survival of WT (n = 79) and STING KO (n = 48) killifish. Indicated p-value represents comparison between STING KO and WT using the log-rank Mantel-Cox test. **c**, Schematic of all fish samples collected from the lifespan experiments in (**a**) and (**b**) for necropsy analysis, with FFPE sections stained with Hematoxylin & Eosin (H&E) and Acid-Fast Bacteria stains. **d**, H&E histological images of the jaw region depicting a healthy and a proliferative non-neoplastic lesion in the lower lip indicated with an asterisk (*). **e**, Heatmap showing the percent occurrence of the proliferative non-neoplastic pathology in 12 tissues analyzed during necropsy in WT and cGAS KO tissues. **f**, H&E histological images depicting a healthy kidney and a kidney displaying degenerative tubular changes. **g**, Percent occurrence of the degenerative pathology in indicated tissues and genotypes. **h**, H&E and Acid-Fast Bacteria-stained histological images of a kidney depicting an inflammatory lesion containing mycobacteria organisms. The outlined area consists of necrosis and mononuclear inflammatory infiltrates. Within the outlined necrotic and inflamed region, arrows point to aggregates of Acid-Fast positive rod-shaped bacteria in the zoomed inset. **i**, Percent occurrence of the inflammatory infectious pathology in indicated tissues and genotypes. Separately, the percent incidence of mycobacteriosis among all tissue is shown. **j**, H&E and Acid-Fast Bacteria-stained histological images of a kidney depicting an inflammatory lesion with no apparent bacteria within it. The inflammatory cell aggregate within the kidney is outlined. Neither positive, nor negative staining acid-fast rod-shaped organisms were detected in these lesions. **k**, Percent occurrence of the inflammatory non-infectious pathology in indicated tissues and genotypes. 60 WT fish and 47 cGAS KO fish were used for necropsy. Bars in images of kidneys = 200 µm. Bars in zoomed in images of kidneys = 10 µm. Bars in jaw images = 500 µm.

We also examined senescence and SASP markers. Surprisingly, in the case of the old kidney, we did not see a strong signature of senescence markers with age (Figure 4i), likely reflecting the small proportion of senescent cells that accumulate within a given tissue during aging [41]. Senescent gene profiles from old cGAS KO kidneys were not very distinct compared to old WT (Figure 4i), with the exception of a handful of genes (NUAK1, CDKN1B, IGFBP1) that appeared less upregulated.

In contrast to the kidney, the aging killifish gut transcriptome showed much less dependence on cGAS (Extended Data Figure 5b, c), which is in line with lack of increased cGAMP activity with age. Pathways significantly regulated during aging were not deregulated in the absence of cGAS, and the dampening effect was less prominent than in the kidney (Extended Data Figure 5d, e). Surprisingly, however, the transcriptomes of old WT guts showed a clearer signature of senescent genes upregulated (CDKN1B/2B, YPEL3, PTBP1) in old age, and lack of cGAS attenuated this regulation (Extended Data Figure 5f).

Unlike cGAS, lack of STING showed hardly any separation of transcriptional profiles with old age when compared to WT in the kidneys (Extended Data Figure 5g, h). However, GSEA showed that proteasome downregulation and complement and coagulation upregulation were blunted in the old STING KO (Figure 4f, Extended Data Figure 5i). In addition, senescent markers (TP63, CDKN1A), SASP regulators (YBX1, MAPK11/14) and SASP components (MIF, IGFP5B, EGF) were downregulated in old STING KO when compared to old WT (Extended Data Figure 5j).

In sum, cGAS modulates the transcriptional landscape during both DNA damage and aging, two conditions that are linked, but the scale of this effect is tissue-specific. STING does not appear to regulate the transcriptional landscape in the kidneys as broadly. Yet, to different degrees, lack of either cGAS or STING attenuates senescent marker and SASP component expression in old age.

### The killifish cGAS/STING pathway affects age-related pathology but not life span

Since killifish cGAS KO and STING KO mitigated senescence in cultured fibroblasts and many age-related transcriptional signatures in the kidney *in vivo*, we sought to investigate the possibility of altered lifespan in our mutants. In particular, we hypothesized that our mutants might live longer than controls due to the amelioration of these age-related phenotypes. To address this, after at least four rounds of backcrossing with WT fish, we generated, expanded, and measured the life span of cGAS and STING KO mutants along with WT fish. Males and females were singly housed, and their life span measured within the same cohort. Demographic analysis showed that cGAS KO fish had fairly similar lifespans to WT fish (Figure 5a, Extended Data Figure 6a-c), contravening our hypothesis. Median and maximum life span of cGAS KO and wild type were comparable and Log rank statistics showed the survival curves were not significantly different (p=0.43). Furthermore, STING KO fish showed a tendency for reduced lifespan (Figure 5b, Extended Data Figure 6a-c), but it did not reach significance (p=0.10).

Hence, we decided to investigate aspects of pathology. In particular, upon death we preserved the fish from the demographic analysis in paraformaldehyde and performed histological necropsy with a certified veterinary pathologist. We categorized the observable disease phenotypes into five categories, namely neoplastic, proliferative non-neoplastic, degenerative, inflammatory infectious and inflammatory non-infectious, and analyzed 12 tissues using 60 WT and 47 cGAS KO preserved fish for necroscopy (Figure 5c). Notably, we found no clear neoplastic lesions in any of the tissues analyzed. Non-neoplastic lesions were detected predominately at the mandibular lip, which was enlarged and contained nodular fibroplasia (Figure 5d), while few fish showed proliferative bronchitis with or without goblet cell hyperplasia. The occurrence of these non-neoplastic lesions was slightly reduced in cGAS KO fish compared to WT (Figure 5e). The most common degenerative lesions were found in the kidney in the form of minimal to mild tubular dilation with or without luminal mineralization (Figure 5f). There, we again observed a tendency for less degenerative phenotype occurrence in the cGAS KO compared to WT kidneys (Figure 5g), consistent with the blunted deregulation that comes with old age in the kidney transcriptome.

Infectious and non-infectious inflammatory lesions were widespread across tissues but most prominent in the kidneys and gonads, specifically the ovaries (Extended Data Figure 6d, e). Infectious lesions often had acid-fast positive rod-shaped bacteria within the lesion, interpreted to be mycobacterial infections (Figure 5h). These infections usually affected multiple organs at once within an individual. Surprisingly, cGAS KO tissues appeared to have a mild reduction in infection occurrence in kidneys, liver, and gonads (Figure 5i), despite this pathway’s involvement in immune defense. Inflammatory non-infectious lesions were classified when significant macrophage infiltration was observed without detectable rod-shaped microorganisms (Figure 5j). These lesions were far less prevalent than infectious lesions but overall, some tissues showed more and others less frequency of occurrence in the cGAS KO (Figure 5k).

Like cGAS KO, STING KO appeared to show a similar reduction in proliferative and degenerative lesion occurrence (Extended Data Figure 6f, g). The reduction of inflammatory lesion occurrence, however, was much more pronounced in the STING mutants, again supporting our observation of a divergent function of STING from cGAS. Infection rates dropped in multiple tissues, especially the gonads (Extended Data Figure 6h), which could be attributed solely to lesions in female ovaries, not male testes. On the contrary, non-infectious inflammatory lesions in the ovary appeared elevated in the STING KO (Extended Data Figure 6i).

It is important to mention that the number of fish used for such histological analyses is not sufficient to definitively determine a protective or detrimental role for each genotype because the differences in odds ratios for most lesions and tissues were small. The only comparison where Fischer’s Exact test revealed a significant change (p=0.034) was the reduction in ovarian infectious lesions in STING KO (3/20 ovaries) compared to WT (15/32 ovaries). Yet the consistent reduction in occurrence of different morbidities across tissues, hints towards an effect of the cGAS/STING pathway beyond ovarian infectious lesions. Most curiously, both cGAS and STING KO appear more susceptible to sterile macrophage infiltration than mycobacterial infections in some tissues.

## Discussion

Inflammaging and immune dysfunction contribute significantly to age-related pathology and disease, and interventions that rebalance these processes can have a profound impact on health and life span [42–44]. The cGAS/STING pathway is a central innate immune signaling pathway that detects cytosolic DNA [7], and has been implicated in senescence cascades induced by DNA damage [11, 19] or mitochondrial dysfunction [12, 20], playing a role in the SASP phenotype [21]. It has been postulated that low grade activation of cGAS/STING over the life course could be a source of chronic inflammation, and in the long term, contribute to aging pathology and organismal demise [3, 6, 7, 45].

In this work, we sought to elucidate the function of the cGAS/STING pathway in the killifish *N. furzeri*, and found that this pathway has conserved roles in senescence, inflammation, and aging. Notably, we found that kcGAS apparently has higher intrinsic enzymatic activity than human cGAS, and can trigger the expression of type I interferon and senescent cell markers in the presence of cytosolic DNA in a heterologous system. Like mammalian counterparts [11, 19], knockout of killifish cGAS and STING mitigates cellular senescence induced by DNA damage in killifish cell culture. In response to irradiation *in vivo*, cGAS KO dampens the transcriptional regulation of genes involved in inflammation, senescence, and the response to DNA damage in young animals, and attenuates changes in the aging transcriptome of old animals relative to WT. Further, both cGAS KO and STING KO fish overall show milder tissue pathology of proliferative non-neoplastic, degenerative, and infectious lesions, consistent with an amelioration of age-related pathology seen in mouse models [3]. Our findings are thus in accord with the hypothesis that inhibition of this conserved pathway could temper age-related inflammation and extend life span.

Yet surprisingly, neither cGAS KO nor STING KO strains are longer lived than wild type. This unexpected result suggests that if there are benefits to these age-related changes, they are offset by other factors that limit lifespan and could arise for various speculative reasons. For example, we found that kidneys from cGAS KO fish showed upregulation of components of the cGAS/STING regulatory cascade, including STING itself, but also a low-grade sterile NF-κB immune signaling, whose chronic activity can be detrimental to tissue physiology [46]. It is also possible that absence of cGAS/STING renders fish more susceptible to viral and bacterial infection. This would certainly be true in the wild [47]. However, laboratory fish aquaria are largely void of fish-infecting viruses [48]. Instead, we generally observed less pathology in the various tissues with decreased cGAS/STING, and somewhat surprisingly, trends to lower levels of infectious pathology rather than non-infectious macrophage infiltration. Perhaps the upregulation of NF-κB signaling or other immune pathways we observed helps ward off such infections. Parallel studies to our own further support that lack of cGAS/STING can increase macrophage infiltration of tissues and even decrease lifespan in murine models [49, 50].

We also observed that killifish cGAS KO fibroblasts in culture had higher levels of γH2AX with or without irradiation, suggesting that cGAS normally prevents intrinsic DNA damage or facilitates clearance *in vitro*. Relatedly, cGAS KO kidney transcriptomes from old fish showed elevated levels of genes involved in DNA repair, as well as DNA replication and cell cycle. Perhaps higher levels of DNA damage alongside relaxation of checkpoints, could lead to cumulative cellular damage and more apoptosis, offsetting any benefit. cGAS has been reported to provide a protective role to genomic DNA through inhibition of transposable elements both at basal levels and post DNA damage [50, 51], whereas other studies suggest that cGAS inhibits DNA repair in cancer cells [52] and bone marrow-differentiated monocytes [53]. Conceivably these disparities could arise from differences in tissue, type of damage, or pathway wiring.

It is possible that the beneficial effects of cGAS/STING inhibition are limited only to specific tissues, which might not have an overall impact on organismal life span. For example, we observed that amelioration of most age-related transcriptional events in the cGAS KO was more pronounced in the kidney, and not as prominent in the intestine. On the other hand, cGAS KO appeared to more greatly attenuate the regulation of senescent genes in the aged intestine. In addition, cGAS and STING KO mostly effected disease occurrence in the kidneys and ovaries, with a smaller effect on other tissues. Different tissues develop varying degrees of DNA damage [54], senescent load [55] and immune activation [56] with age, potentially employing the cGAS/STING pathway to varying degrees. At the same time, senescence and SASP can also offer protective roles [57, 58] and can even stimulate proliferation [21], hence their deregulation might also carry adverse effects. Along similar lines, timing may also be important: lifelong attenuation of such a core signaling cascade might lead to deficient transcriptional responses to stress that counteract the ameliorated ageing transcriptome. Further study of the physiological impact of cGAS/STING across tissues may address these questions in the future.

Although the canonical cytosolic DNA-sensing cGAS/STING pathway involves both cGAS and STING, each protein has functions that are independent of the other. As mentioned above, nuclear cGAS impacts genomic DNA integrity independently of STING [50–52], while STING expression regulates autophagic flux and mitophagy [59]. Studies often investigate the impact of either cGAS or STING assuming they have interchangeable or equal effects, despite the tissue-specific expression level differences between these components [60]. In our study we observed that cGAS and STING often have a similar impact, with varying breadth, while other times their impact was specific to one component. In healthy young kidneys, the lack of cGAS rather than STING led to an increase in basal levels of NF-κB signaling components. During irradiation and ageing, lack of cGAS blunted the regulation of the majority of the most regulated pathways, while the absence of STING had a much more subtle impact, primarily focused around SASP components. Yet, the absence of STING often had a more profound impact on disease occurrence according to our necropsy analysis. Conceivably, some of the observed differences, particularly the activation of low-grade sterile inflammation in young kidneys as well as the attenuation of the transcriptional response to DNA damage-induced senescence and aging, relate to a nuclear role of cGAS and its association with chromatin [61]. Our results underscore the importance of investigating the commonalities and discrepancies in phenotypes observed when inhibiting either cGAS or STING. Understanding their individual roles in cellular and tissue physiology can help in designing safer and more effective clinical applications of cGAS or STING inhibitors, many of which are already undergoing clinical trials [62].

Our study has several limitations as well. Our RNA-seq data was obtained from males and limited to few tissues. Though we carried out demography on males and females, our numbers are not great enough to discern minor sex specific differences. Nevertheless, an overall picture emerges whereby lack of cGAS/STING attenuates cellular senescence in response to DNA damage, and attenuates age-related transcription and pathology, but does not have an effect on organismal life span, suggesting that other processes are limiting for vitality. In the future it will be interesting to see how cGAS and STING individually affect organ specific function and health span at early and later time points of the aging process.

## Material and methods

### Fish husbandry

All experiments were performed on adult (young, aged 7-8 weeks; and old, aged 18-19 weeks) African turquoise killifish *N. furzeri* laboratory strain GRZ-AD. Adult fish were housed singularly in 2.8 L tanks from the second week of life. Water parameters were pH 7–8, kH 3–5 and temperature of 27.5°C. The system was automatically replenished with 10% fresh water daily. Fish were raised in 12 h of light and 12 h of darkness and fed with 10 mg of dry pellet (BioMar INICIO Plus G) and Premium Artemia Coppens twice a day, with total amount of daily food delivered equivalent to 2 – 3% of fish body weight. The first feeding occurred at 8:30, the second at 13:30. For tissue collection, fish were killed by rapid chilling. Tissues were quickly dissected, snap-frozen in liquid nitrogen and stored at −80°C. For irradiation experiments, fish from each genotype were pooled together in a 5 L tank, irradiated with 15 Gy of γ-radiation in a Biobeam 8000 device, and then returned to single housed tanks. Five days post irradiation, fish were sacrificed for tissue collection. Animal experiments were conducted in accordance with relevant guidelines and approved by ‘Landesamt für Natur, Umwelt und Verbraucherschutz Nordrhein-Westfalen’ under license number 2023.A109.

### Generation of cGAS KO and STING KO lines

CRISPR/Cas9 genome editing was performed as described previously [63]. All the single guide RNAs (sgRNAs) were designed based on the CHOP-CHOP web-based tool (https://chopchop.cbu.uib.no). Alt-R S.p. HiFi Cas9 Nuclease and the Alt-R sgRNA were purchased from IDT (see Extended Data table 1 for sgRNA and primer sequences). One-cell-stage embryos were injected with 1 – 2 nl of a solution containing Cas9 enzyme (200 ng µl^−1^), sgRNA (20 ng µl^−1^), KCl (0.2 M) and 1% phenol red. The F0 generation was genotyped by fin-clipping to identify potential founders. Selected founders were then backcrossed with the GRZ-AD strain for at least four generations to remove potential off-target mutations induced by CRISPR editing.

### Establishment of killifish fin fibroblast primary cell cultures

Killifish primary fibroblasts were established as described by Astre et al.[64] with modifications. Specifically, 10-week-old male WT, cGAS KO and STING KO killifish were sacrificed by rapid chilling and their caudal fin excised, washed twice with Dulbecco’s PBS (DPBS, Gibco), and then transferred under a sterile laminar flow hood for subsequent procedures. The fins were washed another two times with sterile DPBS, and then disinfected for 10 min with 25 ppm iodine solution (PVP-I, Sigma-Aldrich) in DPBS. The fins were then washed once with DPBS and the samples further sterilized for 2 h in antibiotic solution with Gentamicin (50ug/ml Gibco) and Primocin® (50ug/ml InvivoGen) in DPBS at room temperature. The fins were then washed again with DPBS and then digested for 20 min in Collagenase P solution (1mg/ml, Merck Millipore) in Leibovitz’s L-15 medium, 2 mM L-glutamine (Gibco). Without washing, the fins were transferred to cell culture plates where they were cut into small pieces using sterile scalpels. The tissues were left on the plate with covers open under the hood for 10 minutes, allowing the digestion solution to evaporate and tissues to adhere to the plate. Plates were then filled with 10 ml L-15 medium supplemented with 15% FBS, Gentamycin (50 ug/ml) and Primocin® (50 ug/ml). The plates were carefully washed with DPBS so that the tissues did not detach, and media was refreshed every 4 days. When egress of cells from the tissues became visible, the tissues were removed while the cells were then detached with Trypsin-EDTA 0.05% (Gibco) and passaged onto a new plate. For the first 5 passages, cells were grown in media containing Gentamycin and Primocin (50 ug/ml each), and thereafter grown without antibiotics.

### Cell culture maintenance

cGAS KO and STING KO human THP1 monocytes were purchased from Invivogen (THP1-Dual™ KO-cGAS Cells and THP1-Dual™ KO-STING Cells). THP1 cells were cultured under 5% CO_2_ at 37°C in RPMI 1640 (Gibco), 4.5 g/L D-Glucose, 2.383 g/L HEPES, 2 mM L-glutamine, 1.5 g/L Sodium Bicarbonate, 110 mg/L Sodium Pyruvate, 50 µM 2-Mercaptoethanol (Gibco), 20% (v/v) heat-inactivated fetal bovine serum (FBS; 30 min at 56°C for heat inactivation). Killifish fin fibroblasts were cultured in a humidified 28°C incubator with no addition of CO_2_ in Leibovitz’s L-15 medium (Gibco), 2 mM L-glutamine, 15% (v/v) fetal bovine serum.

Both human and killifish cells were repeatedly tested for mycoplasma infection using Eurofins Genomics Mycoplasmacheck Service testing for all mycoplasma species listed in the European Pharmacopoeia 2.6.7.

### Primary cell culture irradiation and senescence-associated β-galactosidase staining

Killifish primary cells were irradiated with 10 Gy of γ-radiation in a Biobeam 8000 device 24h after the last passage, while still in the medium-filled plate. For senescence-associated β-galactosidase staining, cells were fixed and stained nine days post-irradiation using the Senescence β-Galactosidase Staining Kit (Cell Signaling) according to the manufacturer’s instructions. Images of the cells were taken using an EVOS FL Auto 2 microscope and analyzed with the EVOS FL Auto 2 Cell Imaging System software.

### Plasmids

All plasmids were generated using the pSBbi-GN backbone, designed by Eric Kowarz [65] (Addgene ID #60517). The kcGAS and kSTING genes were amplified from cDNA obtained from the killifish primary cell cultures using primers with overhangs matching the Gibson Assembly regions on pSBbi-GN (see Extended Data Table 1 for primer sequences). Similarly, the hcGAS and hSTING genes were amplified from WT THP1 (InvivoGen) cDNA. The pSBbi-GN vector was linearized with SfiI (NEB), and the linear plasmid was combined with the amplified cDNAs for Gibson Assembly (NEBuilder® HiFi DNA Assembly, NEB). All cloning products used in the experiments have been submitted to Addgene (kcGAS: #225274, hcGAS #225275, kcGAS KO #225276, kSTING #225277, hSTING #225278, kSTING KO #225279).

### Cell culture transfections

THP1 cells were transfected using GeneXPlus (ATCC) according to the manufacturer’s instructions. 48 h after transfection, the cells were split in three equal parts. One part was used for 2’,3’-cGAMP measurement, one part was used for RNA extraction and qPCRs and one part was used for flow cytometry to determine transfection efficiency.

### LC–MS-based targeted metabolomic measurements of cyclic nucleotides

The method was adapted from our previous work [31]. Briefly, cells were extracted using a two-phase extraction method with a mixture of DW:MeOH:CHCl3 (1:2:2). After centrifugation, the separated upper and lower layers were dried under a vacuum evaporator and stored at –80 °C until further analysis. The protein content in the cell pellet was then used for normalization, determined by the PierceTM BCA protein assay kit (Thermo Fisher Scientific).

Compounds were separated using a reversed-phase column (HSS T3 column 1.8 μm, 100 mm × 2.1 mm, Waters) at 40 °C by the Vanquish UHPLC system (ThermoFisher Scientific GmbH, Bremen, Germany) coupled to a triple quadrupole mass spectrometer (TSQ Altis, ThermoFisher Scientific GmbH, Bremen, Germany).

The mobile phases consisted of 1% formic acid in H_2_O (A) and 1% formic acid in ACN (B) with a flow rate of 0.3 mL/min. The gradient was set as follows: 1% B was maintained for 1 min, then ramped up to 15% B over 3 min and reached 50% in an additional 1 min. At min 7, it reached 70% B and was maintained for 1.5 min. The gradient then quickly decreased to 1% B and re-equilibrated for 3 min.

2′3′-cGAMP (m/z 675.67) was quantified using a fragment ion at m/z 476.01 and validated by an ion at m/z 505.9. The signal was calculated by dividing the peak area by an internal standard peak area (adenosine-13C10,15N5-5′-monophosphate, m/z 363.12 → m/z 146.08) and further normalized to protein concentration. Data were analyzed using Skyline Version 22.2.0.527.

### RNA extraction and qPCR analysis

Killifish tissues were ground in a liquid nitrogen filled mortar using a pestle until completely pulverized. The powder was then thawed in RLT buffer (QIAGEN) with 1% β-ME v/v on ice. Samples were then centrifuged at 10.000 g for 10 min at 4°C. The supernatant was collected for subsequent RNA extraction.

As non-adherent cells, THP1 cells were collected by centrifugation at 500 g for 4 min at room temperature. The pellet was washed once with DPBS and then snap frozen in liquid nitrogen. The frozen pellets were thawed in RLT buffer (QIAGEN) with 1% β-ME v/v on ice for subsequent RNA extraction.

Nine days post irradiation, killifish fibroblasts on culture dishes were washed with PBS and then incubated with RLT buffer (QIAGEN) with 1% β-ME v/v on ice for 5 min, before the cells were scraped and used for subsequent RNA extraction.

For all above samples in RLT, RNA extraction was performed using the RNeasy Mini kit (QIAGEN) according to the manufacturer’s instructions. The optional DNase step was always performed using the RNase-Free DNase Set kit (QIAGEN) according to the manufacturer’s instructions. The concentration and purity of the RNA were measured by NanoDrop. cDNA was generated using iScript (Bio-Rad). qPCR with reverse transcription was performed with Power SYBR Green (Applied Biosystems) on a ViiA 7 Real-Time PCR System (Applied Biosystems). Four technical replicates were averaged for each sample per primer reaction. GAPDH and EIF3C were used as internal controls for killifish samples, GAPDH was used for human THP1 cells (Extended Data table 1 provides primer sequences).

### Flow cytometry

THP1 cells were pelleted by centrifugation at 500 g for 4 min, then washed once with DPBS (Gibco) and pelleted again the same way. Cell pellets were resuspended in cold 2% (v/v) FBS in DPBS and passed through a 35 µm nylon mesh cell strainer to obtain single cells. 0.1 µg/mL DAPI was added to each sample and at least 10.000 cells were then immediately recorded using a BD FACSCanto II flow cytometer. Data were analyzed using FloJo software. Briefly, cells were gated based on forward and side scatter and doublets were excluded. Gates for GFP positive cells were adjusted by comparison to untransfected THP1 cells. Gates for DAPI positive cells were adjusted by comparison to samples without DAPI. Compensation between DAPI and GFP was made using samples without DAPI and/or GFP.

### Immunofluorescence

Killifish cells were plated on glass slide chambers (Nunc™ Lab-Tek™ II) where they were allowed to attach for 24 h. Then starting with the 24 h timepoint, the chambers were irradiated with 10 Gy of γ-radiation in a Biobeam 8000 device and then placed back in a humidified 28°C incubator without CO_2_ supplementation. After all timepoints were completed (8 h, 4 h, 2 h and non-irradiated 0 h), the slides were processed together at room temperature unless otherwise specified. They were washed with DPBS (Gibco) once and then fixed in freshly prepared 4% (w/v) paraformaldehyde (Sigma-Aldrich) in PBS with adjusted pH 7.2 for 15 min. They were then washed three times with DPBS and permeabilized with 0.5% (v/v) Triton-X100 (Sigma-Aldrich) in DPBS for 10 min. They were washed again 3 times with DPBS and then blocked for 1 h in 8% (v/v) FBS, 0.3% (v/v) Triton-X100 in DPBS. After blocking, primary antibody for γH2AX (#9718 Cell Signaling) was used at 1:200 dilution in 5% (v/v) FBS, 0.1% (v/v) Triton-X100 in DPBS and incubated at 4°C overnight. The slides were washed three times with DPBS and then incubated for 1 h with fluorescence-conjugated secondary antibody (#A21206, Invitrogen) diluted 1:1000 in 5% (v/v) FBS, 0.1% (v/v) Triton-X100 in DPBS. The secondary antibody solution was washed off the slides 3 times and the slides were mounted using Fluoromount-G™ with DAPI (Invitrogen). Images were taken under a Leica DMI6000 B microscope and analyzed with FIJI. Specifically, a stringent intensity threshold was set such that only intense and separate puncta were counted. Due to blinding, images with distinct puncta were chosen randomly to set the threshold, which was then applied across all images. After thresholding, only objects with size 1-50 pixels were counted per cell in order to avoid clumps. At least 50 cells per slide were imaged.

### RNAseq analysis

Kidneys and guts (stomach and intestine) from cGAS KO as well as kidneys from STING KO fish were collected for RNAseq analysis. The young irradiated and non-irradiated kidneys from STING KO fish were processed in one batch, while all other samples were processed in another. Both WT and KO mutants derived from offspring of heterozygous parents to ensure similar genetic backgrounds. To reduce circadian variability, fish in each batch were killed all at once within 2 h in the early afternoon. Collected tissues were snap-frozen in liquid nitrogen and stored at −80°C. RNA extraction of all samples in each batch was performed at the same time.

Libraries were prepared from 500 ng total RNA. ERCC RNA Spike-In Mix 1 (Thermo Fischer) was added to the samples before library preparation. Enzymatic depletion of ribosomal RNA with the Ribo-Zero Gold rRNA Depletion kit (Illumina) was followed by library preparation with the TruSeq Stranded Total RNA sample preparation kit (Illumina). The depleted RNA was fragmented and reverse transcribed with random hexamer primers, second strand synthesis with dUTPs was followed by A-tailing, adapter ligation and library amplification (15 cycles). Next, library validation and quantification (Agilent Tape Station) were performed, followed by pooling of equimolar amounts of libraries. The library pools were then quantified using the Peqlab KAPA Library Quantification Kit and the Applied Biosystems 7900HT Sequence Detection System and sequenced on an Illumina NovaSeq6000 sequencing instrument with a PE100 protocol aiming for 50 million clusters per sample.

After removal of residual rRNA and tRNA reads, remaining reads were pseudo aligned to the reference genome (Nfu_20140520) using Kallisto (v.0.45.0) [66] and RSeQC/4.0.0 was used to identify mapping strand [67]. A strand was identified by having more than 60% of reads mapped to it. Cases with less than 60% of reads in each strand were defined as unstranded. Genes with fewer than ten overall reads were removed. After normalization of read counts making use of the standard median ratio for estimation of size factors, pairwise differential gene expression analysis was performed using DESeq2 (v.1.24.0) [68]. The log2 fold changes were shrunk using approximate posterior estimation for GLM coefficients. Gene Set Enrichment Analysis (GSEA) [69, 70] and Principal Component Analysis (PCA) were performed for all comparisons using Flaski (https://flaski.age.mpg.de/).

### Tissue fixation, H&E staining, and Acid-Fast Bacteria staining

Fish within the lifespan cohorts were monitored 4 times a day to check for viability. Dead fish were collected, their visceral cavity was opened, and then they were placed in 10% Formalin solution at room temperature. Fish with severe post-mortem autolysis typically due to 16-24 hr between death and detection, were not included in the study. The fish were kept in Formalin until the completion of the lifespan experiment. 141 samples were then submitted for processing and histopathologic evaluation by IDEXX BioAnalytics. Briefly, all fish were paraffin embedded and 4 sections were taken from each fish. After deparaffinization the sections were stained for either H&E or Acid-Fast Bacteria (AFB) staining. The presence or absence of neoplastic, proliferative but non-neoplastic, degenerative, non-infectious inflammatory, and infectious inflammatory lesions were recorded for a routine tissue list including the brain, epidermis, gills, heart, digestive tract (headgut, foregut, midgut, hindgut), kidney, liver, gonads (ovary or testes), skeletal muscle, spinal cord, swim bladder, and vertebral column.

### Lifespan analysis

For both cGAS and STING mutants, P0 heterozygous fish were bred to produce the F1 KO fish that were the parents of all F2 fish used in the lifespan. All fish used in the lifespan derived from eggs collected within the span of 10 days. After hatching, larvae were housed together (seven larvae per 1.1 L tank), until they reached 2 weeks of age. Then they were single-housed in 2.8 L tanks for the remaining lifespan. Fish mortality was scored starting at the sixth week, when full sexual maturation was reached. By that time, no fish had detectable abnormalities in body size or swimming and as such no fish were censored from the study. Senescent fish (> 28 weeks) that showed visible signs of stress, lethargy, anorexia, and abnormal swimming behaviors were sacrificed for humane reasons. The age of fish sacrificed for such reasons was recorded as the estimate time of natural death. Survival curves were calculated using the Kaplan–Meier estimator. Statistical significance was calculated by the Mantel Cox log-rank test.

## Data availability

All raw RNAseq data are available in the SRA database under BioProject ID PRJNA1154547. All data supporting the findings of this study are available from the corresponding author upon request.

## Acknowledgements

We would like to especially thank the MPI-AGE fish facility for fish maintenance and husbandry, the FACS and Imaging, and Bioinformatics cores for their services, members of the Antebi lab for helpful discussions, Ray Laboy for performing the Alphafold 3 analysis, Sarah Kreuz for critical reading of the manuscript and the Max Planck Gesellschaft for funding. Eugen Ballhysa received funding from the Cologne Graduate School of Ageing Research.

## Contributions of Authors

A.A. and R.R. conceived the study. E.B., R.R. and A.A. conceived and designed the experiments. E.B. performed all the experiments with input from R.R. N.H. offered technical assistance. LC-MS analysis was performed by T.T.M.N. Histopathology was performed by J.B. B.F. and E.H. helped with cloning and establishing primary killifish fibroblasts. J.S. helped analyze flow cytometry data. E.B. and A.A. wrote the manuscript. All authors reviewed and approved the manuscript.

## Extended Data

**Extended Data Figure 1.**
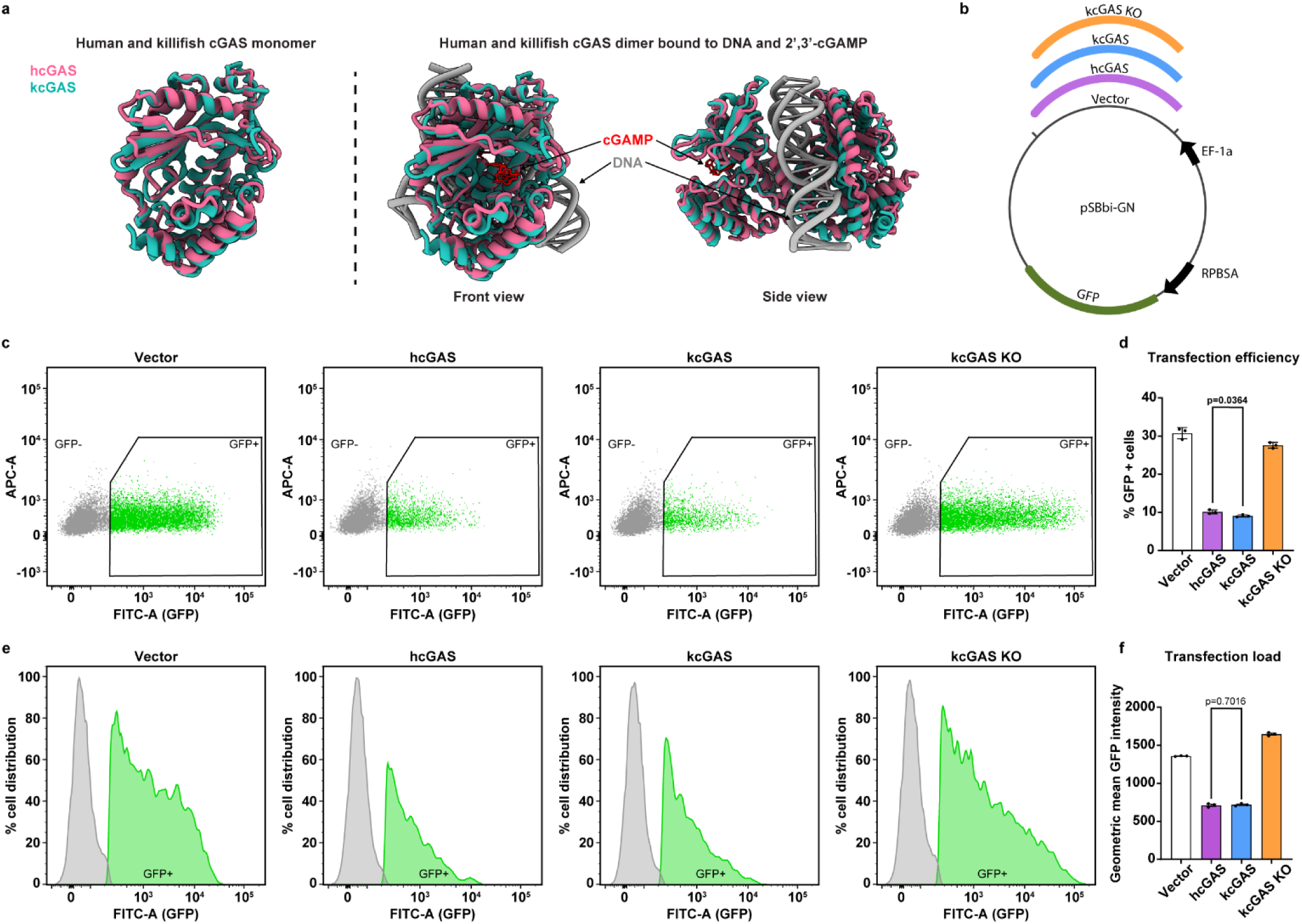
Killifish and human cGAS have similar tertiary structures and affect THP1 cell transfection efficiency. **a**, Structural alignment of the human cGAS (PDB 5VDO) with the AlphaFold 3-predicted model of killifish cGAS. Left, monomers of killifish (turquoise) and human (purple) cGAS proteins are shown in superposition. Right, two strands of DNA are flanked by kcGAS and hcGAS in superposition in the dimeric conformation. Also shown bound is the enzymatic product 2’,3’-cGAMP. **b**, Schematic of the expression vector used for transfecting hcGAS, kcGAS and kcGAS KO to THP1 cells. Genes of interest are constitutively expressed under the control of the EF-1a promoter, while in parallel, GFP is constitutively expressed under control of the RPBSA promoter. Plasmid maps and sequences can be found in Extended Data table 1. **c**, Representative dot plots from flow cytometry of alive THP1 cells after transfection with plasmids carrying the indicated cGAS genes. The GFP intensity on the x-axis (FITC-A) is plotted against background noise intensity of an empty gate (APC-A). Cells within the GFP+ gate are considered successfully transfected. **d**, Quantification of transfected cGAS KO THP1 cell percentages falling within GFP+ gate as shown in (**c**). Student’s t-test was used to compare hcGAS and kcGAS means. **e**, Histograms depicting the distribution of GFP fluorescence intensity within cell populations of transfected THP1 cells. The GFP intensity on the x-axis (FITC-A) is plotted against the percent distribution of cells at each intensity level. The grey peak represents alive GFP-cells, while the green peak represents alive GFP+ cells. **f**, Geometric means of GFP intensities of the GFP+ cells shown in (**e**). Student’s t-test was used to compare hcGAS and kcGAS geometric means. Flow cytometry data in **c**-**e** were obtained from the same samples as those used in Figure 1d, **e** and are thus representative of the transfection efficiency that comes with those data. Experiments were performed in 3 independent biological replicates with n = 3 plates each time. Each panel shows one of three independent experiments.

**Extended Data Figure 2.**
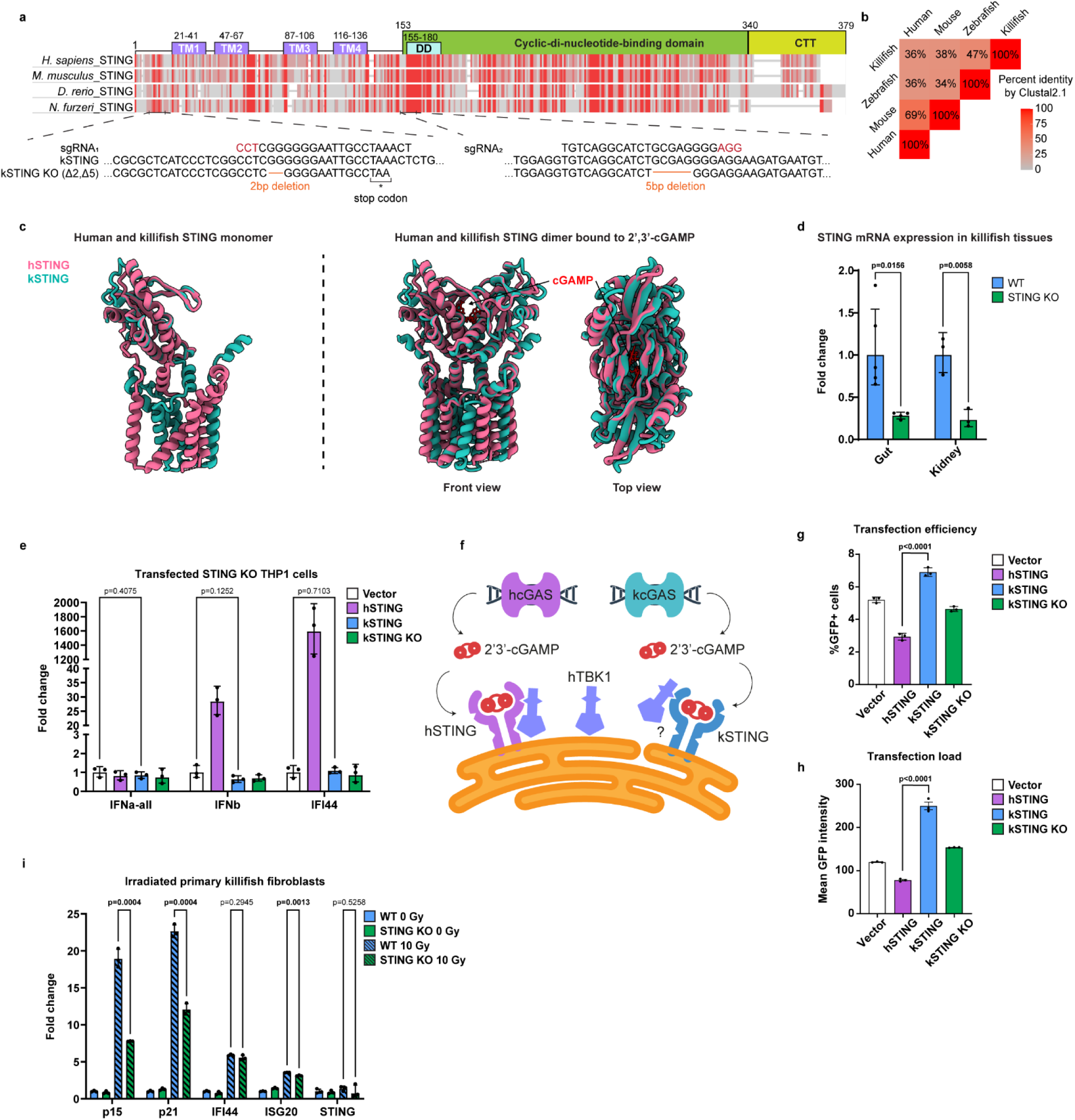
STING structure and function are conserved from teleosts to humans. **a**, Multiple sequence amino acid alignment of STING1 (STING) with orthologues from other species using constraint-based multiple sequence alignment tool (COBALT). Red= identical amino acids, grey= non-identical. Above the alignment are shown the four transmembrane domains (TM1-4), the dimerization domain (DD), the cyclic-di-nucleotide-binding domain and the C-terminal tail (CTT). Below the alignment is a schematic of the CRISPR-Cas9-generated mutated region of killifish STING and the used sgRNAs. **b**, Triangle heatmap showing percent identity between aligned STING genes of different organisms using ClustalW. **c**, Structural alignment of the human STING (PDB 8FLM) using the AlphaFold 3-predicted model of killifish STING. Left, monomers of killifish (turquoise) and human (purple) STING proteins are shown in superposition. Right, the conformation of hSTING and kSTING dimers bound to 2’,3’-cGAMP is shown. **d**, Expression of STING mRNA measured using qPCR in different tissues of WT and STING KO mutant killifish. Each dot represents data from one fish. n ≥ 3 per tissue and genotype. Statistical analysis was done with Student’s t-test using the normalized expression values of STING to GAPDH. **e**, Expression of interferon and interferon-stimulated genes by qPCR from cell extracts of STING KO THP1 cells transfected with the indicated STING genes. Statistical analysis was done with Student’s t-test using the normalized expression values to GAPDH. **f**, Schematic depicting human and killifish cGAS/STING signaling. Both human and killifish cGAS produce the same secondary messenger 2’,3’-cGAMP. The illustration also shows the potential lack of binding affinity of human TBK1 protein to kSTING. **g**, Quantifications of transfected STING KO THP1 cell percentages after flow cytometric analysis of cells falling within the GFP+ gate. Gating strategy was similar to the one shown in Extended Data Figure 1C, but baseline adjusted to levels of the control untransfected STING KO THP1 cells. Student’s t-test was used to compare hSTING and kSTING means. **h**, Geometric means of GFP intensities of the GFP+ THP1 cells from (**g**). Student’s t-test was used to compare hSTING and kSTING means. **i**, qPCR measurement of senescence and interferon-stimulated genes in irradiated WT and STING KO primary fibroblasts. Statistical analysis was done with Student’s t-test using the normalized expression values to EIF3c. STING KO THP1 cell transfections in **e**-**h** were performed in 2 independent biological replicates with n = 3 plates each time. qPCR on irradiated primary STING KO fibroblasts in **i** was performed in 3 independent biological replicates with n = 3 plates each time. Each panel shows one of three independent experiments.

**Extended Data Figure 3.**
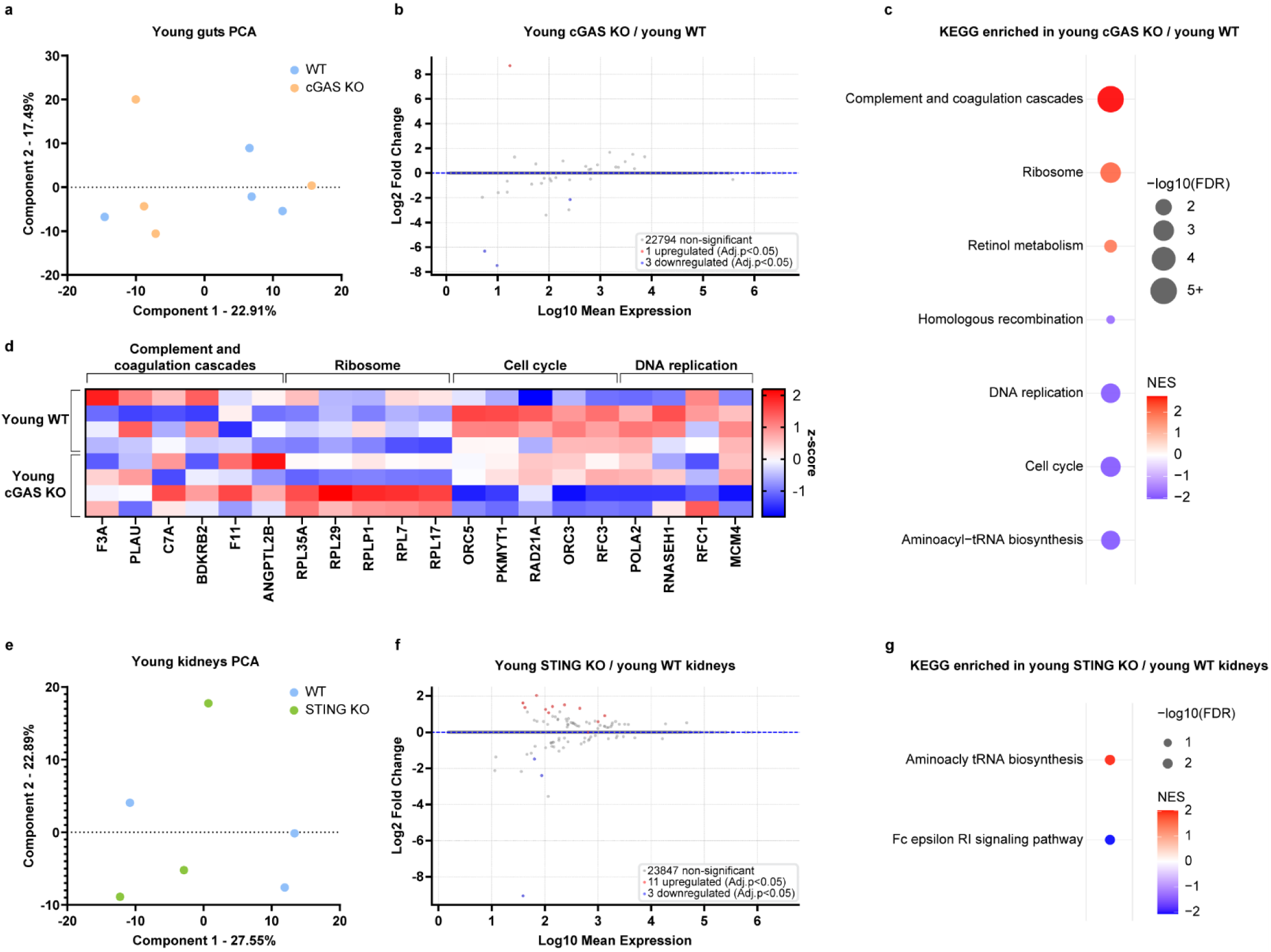
Lack of cGAS moderately impacts the gut transcriptome, while lack of STING marginally impacts the kidney transcriptome of young fish. **a**, PCA plot of WT and cGAS KO transcriptomes from young (8 weeks) killifish guts. **b**, MA plot showing differential mRNA expression comparing cGAS KO to WT young guts. All genes with adjusted p-value < 0.05 are considered significant and marked as upregulated (red) or downregulated (blue). **c**, GSEA comparing transcriptomes of cGAS KO to WT young guts. All KEGG pathways significantly upregulated and downregulated in cGAS KO are shown (FDR < 0.05). **d**, Heatmap of z-score normalized expression values of the genes with the highest positive and lowest negative rank metric scores from representative GSEA pathways shown in (**c**). **e**, PCA plot of WT and STING KO transcriptomes from young (8 weeks) killifish kidneys. **f**, MA plot showing differential mRNA expression comparing STING KO to WT young kidneys. All genes with adjusted p-value < 0.05 are considered, upregulated (red), downregulated (blue). **g**, GSEA comparing transcriptomes of STING KO and WT young kidneys. All KEGG pathways significantly upregulated and downregulated in STING KO are shown (FDR < 0.05). NES: Normalized enrichment score, FDR: False discovery rate. n = 4 fish per genotype.

**Extended Data Figure 4.**
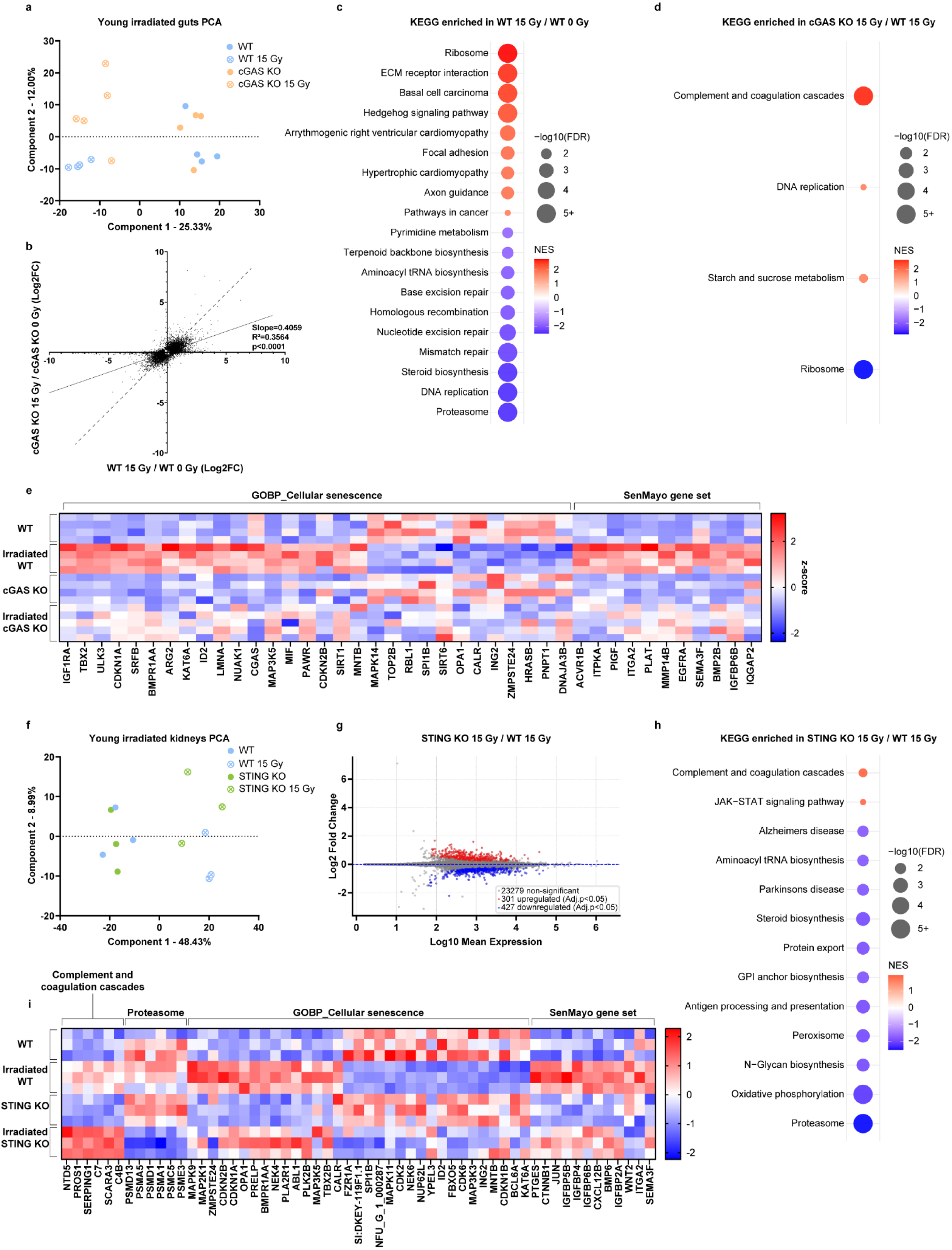
cGAS and STING modulate DNA damage induced transcriptional changes in guts and kidneys. **a**, PCA plot of WT and cGAS KO transcriptomes from guts of irradiated and non-irradiated young (8 weeks) killifish. **b**, Linear regression analysis of the Log2(Fold change) of all genes significantly changed in expression after irradiation in WT and cGAS KO guts, showing a slope significantly < 1. Dashed line with slope = 1 is shown for comparison. **c**, GSEA of guts from irradiated compared to non-irradiated WT fish. The top KEGG pathways upregulated and downregulated in irradiated WT are shown. **d**, GSEA of guts from irradiated cGAS KO compared to WT killifish. The top KEGG pathways upregulated and downregulated in cGAS KO are shown. **e**, Heatmap of z-score normalized expression values of genes with the highest positive and lowest negative rank metric scores from GSEA when comparing guts from irradiated to non-irradiated WT fish. The gene sets investigated were the Sen_Mayo gene set and the GOBP_Cellular Senescence gene set. **f**, PCA plot of WT and STING KO transcriptomes of kidneys from irradiated and non-irradiated young (8 weeks) killifish. **g**, MA plot showing differential mRNA expression comparing kidneys from young irradiated STING KO to kidneys from irradiated WT killifish. All genes with adjusted p-value < 0.05 are considered significant, upregulated (red), downregulated (blue). **h**, GSEA of kidneys from irradiated STING KO compared to WT killifish. The top KEGG pathways upregulated and downregulated in STING KO are shown. **i**, Heatmap of z-score normalized expression values of genes with the highest positive and lowest negative rank metric scores from GSEA when comparing guts from irradiated to non-irradiated WT fish. Genes shown derive from representative pathways from (**h**) as well as Sen_Mayo gene set and the GOBP_Cellular Senescence gene set.

**Extended Data Figure 5.**
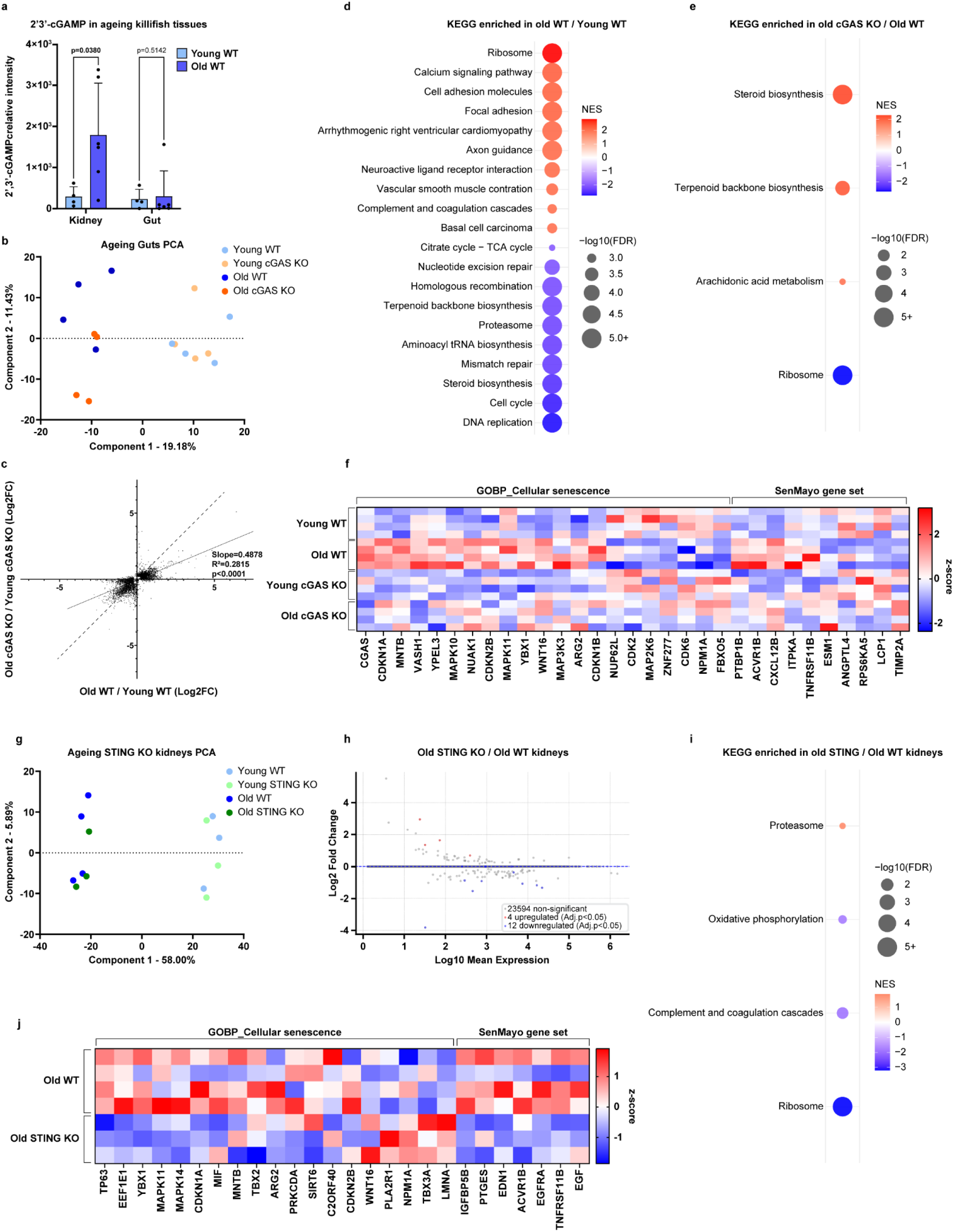
cGAS and STING modulate senescent transcriptional signatures during ageing in killifish kidneys and guts. **a**, LC–MS-based targeted metabolomics of 2’,3’-cGAMP extracted from young (7 weeks, n = 4) and old (18 weeks, n = 6) WT kindeys and guts. Statistical analysis was performed using Mann–Whitney U test. **b**, PCA plot of WT and cGAS KO transcriptomes from young (8 weeks) and old (18 weeks) killifish guts. For each condition n = 4. **c**, Linear regression analysis of the Log2(Fold change) of all genes significantly changed in expression during ageing in WT and cGAS KO guts, showing a slope significantly < 1. Dashed line with slope = 1 is shown for comparison. **d**, GSEA comparing old to young WT guts. The top KEGG pathways upregulated and downregulated in old WT guts are shown. **e**, GSEA comparing old cGAS KO to WT guts. All KEGG pathways upregulated and downregulated in cGAS KO are shown. **f**, Heatmap of z-score normalized expression values of genes with the highest positive and lowest negative rank metric scores from GSEA when comparing old to young WT guts. The gene sets investigated were the Sen_Mayo gene set and the GOBP_Cellular Senescence gene set. **g**, PCA plot of WT and STING KO transcriptomes from young (9 weeks) and old (18 weeks) killifish kidneys. For old WT n = 4, for all other conditions n = 3. **h**, MA plot showing differential mRNA expression comparing old STING KO to old WT kidneys. All genes with adjusted p-value < 0.05 are considered significant, upregulated (red), downregulated (blue). **i**, GSEA comparing old STING KO to WT kidneys. All KEGG pathways upregulated and downregulated in STING KO are shown. **j**, Heatmap of z-score normalized expression values of genes with the highest positive and lowest negative rank metric scores from GSEA when comparing old STING KO to WT kidneys. The gene sets investigated were the Sen_Mayo gene set and the GOBP_Cellular Senescence gene set.

**Extended Data Figure 6.**
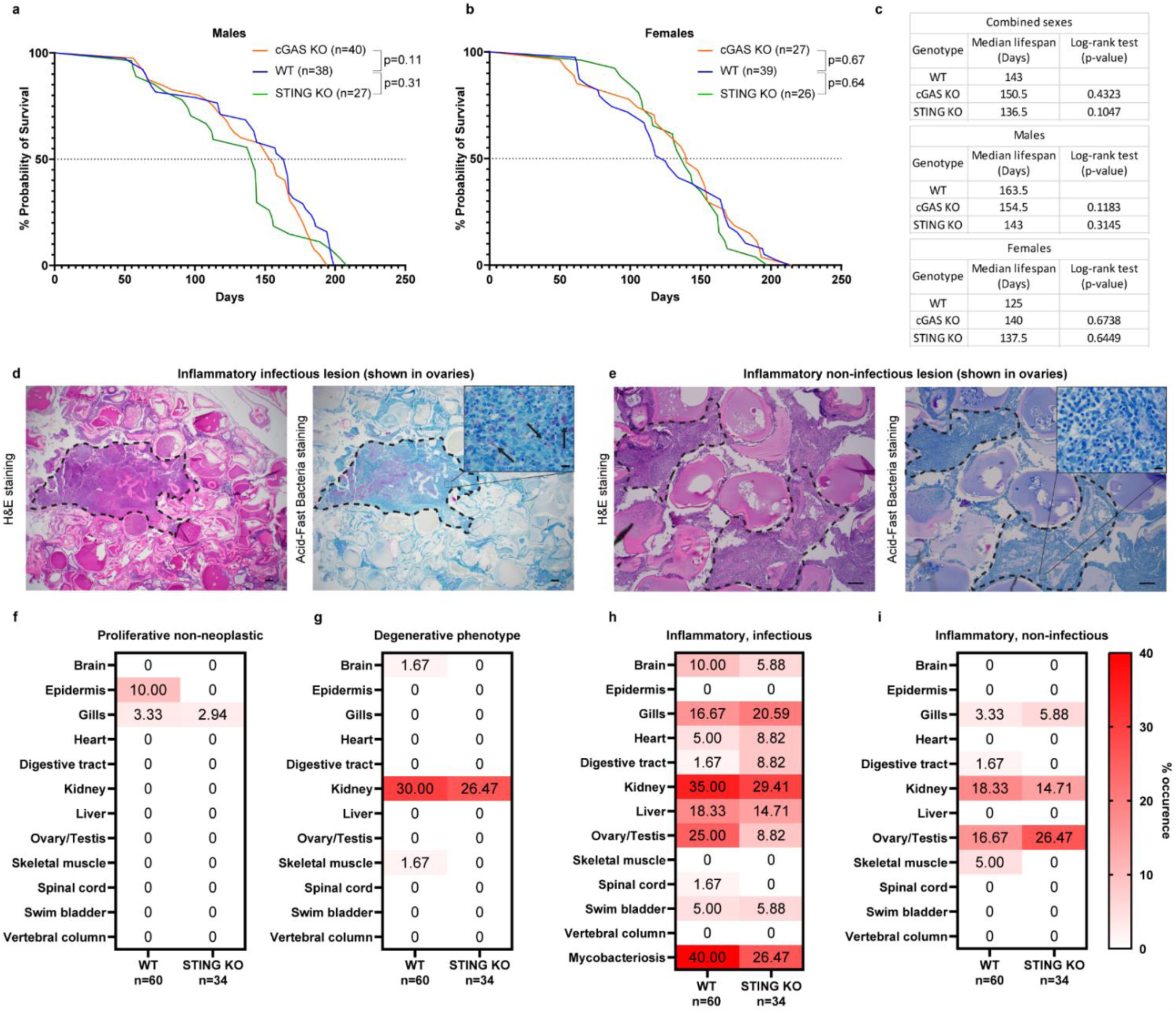
The killifish cGAS/STING pathway contributes to degenerative and inflammatory disease occurrence but not lifespan in individual sexes. **a**, Kaplan-Meier curve showing the survival of WT (n = 38), cGAS KO (n = 40) and STING KO (n = 27) male killifish. Indicated p-values represent statistical comparisons between WT and cGAS KO or STING KO, respectively, using the log-rank Mantel-Cox test. **b**, Kaplan-Meier curve showing the survival of WT (n = 39), cGAS KO (n = 27) and STING KO (n = 26) female killifish. Indicated p-value represent statistical comparisons between WT and cGAS KO or STING KO, respectively, using the log-rank Mantel-Cox test. **c**, Tables summarizing the median lifespans (in days) of WT, cGAS KO and STING KO killifish. Both individual sexes and combined sexes are shown separately. Log-rank tests were performed comparing WT to cGAS or STING KO fish, respectively, and the p-values are shown for each genotype. **d**, H&E and Acid-Fast Bacteria-stained histological images of a killifish ovary depicting an inflammatory lesion containing mycobacteria organisms. The inflammatory cell aggregate within the necrotic ovarian region is outlined. Within the necrotic and inflamed region, arrows point to clusters of Acid-Fast positive rod-shaped bacteria, shown in the zoomed inset. **e**, H&E and Acid-Fast Bacteria-stained histological images of a killifish ovary depicting an inflammatory lesion containing no apparent bacteria. The inflammatory cell aggregate within the ovary is outlined. Neither positive, nor negative Acid-Fast rod-shaped organisms were detected in these lesions. **f**, Heatmap showing the percent occurrence of proliferative non-neoplastic pathology in 12 tissues as analyzed during necropsy in WT and STING KO tissues. **g**, Percent occurrence of the degenerative pathology in indicated tissues and genotypes. **h**, Percent occurrence of the inflammatory infectious pathology in indicated tissues and genotypes. Separately, the percent incidence of mycobacteriosis among all tissues is shown. **i**, Percent occurrence of inflammatory non-infectious pathology in indicated tissues and genotypes. 60 WT fish and 34 STING KO fish were used for necropsy. Bars in images of ovaries = 200 µm. Bars in inset images of ovaries = 10 µm.

## References

1. Franceschi, C., et al., Inflamm-aging. An evolutionary perspective on immunosenescence. Ann N Y Acad Sci, 2000. 908: p. 244–54.

2. Yu, H., et al., Role of the cGAS-STING Pathway in Aging-related Endothelial Dysfunction. Aging Dis, 2022. 13(6): p. 1901–1918.

3. Gulen, M.F., et al., cGAS-STING drives ageing-related inflammation and neurodegeneration. Nature, 2023. 620(7973): p. 374–380.

4. Baruch, K., et al., Aging. Aging-induced type I interferon response at the choroid plexus negatively affects brain function. Science, 2014. 346(6205): p. 89–93.

5. Samson, N. and A. Ablasser, The cGAS-STING pathway and cancer. Nat Cancer, 2022. 3(12): p. 1452–1463.

6. Franceschi, C., et al., Inflammaging: a new immune-metabolic viewpoint for age-related diseases. Nat Rev Endocrinol, 2018. 14(10): p. 576–590.

7. Decout, A., et al., The cGAS-STING pathway as a therapeutic target in inflammatory diseases. Nat Rev Immunol, 2021. 21(9): p. 548–569.

8. Civril, F., et al., Structural mechanism of cytosolic DNA sensing by cGAS. Nature, 2013. 498(7454): p. 332–7.

9. Gluck, S., et al., Innate immune sensing of cytosolic chromatin fragments through cGAS promotes senescence. Nat Cell Biol, 2017. 19(9): p. 1061–1070.

10. Mackenzie, K.J., et al., cGAS surveillance of micronuclei links genome instability to innate immunity. Nature, 2017. 548(7668): p. 461–465.

11. Yang, H., et al., cGAS is essential for cellular senescence. Proceedings of the National Academy of Sciences of the United States of America, 2017. 114(23): p. E4612–E4620.

12. West, A.P., et al., Mitochondrial DNA stress primes the antiviral innate immune response. Nature, 2015. 520(7548): p. 553–7.

13. Sprenger, H.G., et al., Cellular pyrimidine imbalance triggers mitochondrial DNA-dependent innate immunity. Nat Metab, 2021. 3(5): p. 636–650.

14. Du, Y., et al., Function and regulation of cGAS-STING signaling in infectious diseases. Front Immunol, 2023. 14: p. 1130423.

15. Sun, L., et al., Cyclic GMP-AMP synthase is a cytosolic DNA sensor that activates the type I interferon pathway. Science, 2013. 339(6121): p. 786–91.

16. Wu, J., et al., Cyclic GMP-AMP is an endogenous second messenger in innate immune signaling by cytosolic DNA. Science, 2013. 339(6121): p. 826–30.

17. Ablasser, A., et al., cGAS produces a 2’-5’-linked cyclic dinucleotide second messenger that activates STING. Nature, 2013. 498(7454): p. 380–4.

18. Zheng, W., et al., The Role of cGAS-STING in Age-Related Diseases from Mechanisms to Therapies. Aging Dis, 2023. 14(4): p. 1145–1165.

19. Dou, Z.X., et al., Cytoplasmic chromatin triggers inflammation in senescence and cancer. Nature, 2017. 550(7676): p. 402–406.

20. Victorelli, S., et al., Author Correction: Apoptotic stress causes mtDNA release during senescence and drives the SASP. Nature, 2024. 625(7995): p. E15.

21. Coppe, J.P., et al., The senescence-associated secretory phenotype: the dark side of tumor suppression. Annu Rev Pathol, 2010. 5: p. 99–118.

22. Khosla, S., et al., The role of cellular senescence in ageing and endocrine disease. Nat Rev Endocrinol, 2020. 16(5): p. 263–275.

23. Platzer, M. and C. Englert, Nothobranchius furzeri: A Model for Aging Research and More. Trends Genet, 2016. 32(9): p. 543–552.

24. Ge, R., et al., Conservation of the STING-Mediated Cytosolic DNA Sensing Pathway in Zebrafish. J Virol, 2015. 89(15): p. 7696–706.

25. de Oliveira Mann, C.C., et al., Modular Architecture of the STING C-Terminal Tail Allows Interferon and NF-kappaB Signaling Adaptation. Cell Rep, 2019. 27(4): p. 1165–1175 e5.

26. Sellaththurai, S., et al., CRISPR/Cas9-Induced Knockout of Sting Increases Susceptibility of Zebrafish to Bacterial Infection. Biomolecules, 2023. 13(2).

27. Liu, Z.F., et al., Characterization of cGAS homologs in innate and adaptive mucosal immunities in zebrafish gives evolutionary insights into cGAS-STING pathway. FASEB J, 2020. 34(6): p. 7786–7809.

28. Valdesalici, S. and A. Cellerino, Extremely short lifespan in the annual fish Nothobranchius furzeri. Proc Biol Sci, 2003. 270 Suppl 2(Suppl 2): p. S189–91.

29. Kim, Y., H.G. Nam, and D.R. Valenzano, The short-lived African turquoise killifish: an emerging experimental model for ageing. Dis Model Mech, 2016. 9(2): p. 115–29.

30. Boos, F., J. Chen, and A. Brunet, The African Turquoise Killifish: A Scalable Vertebrate Model for Aging and Other Complex Phenotypes. Cold Spring Harb Protoc, 2024. 2024(3): p. 107737.

31. Annibal, A., et al., Mass spectrometric characterization of cyclic dinucleotides (CDNs) in vivo. Anal Bioanal Chem, 2021. 413(26): p. 6457–6468.

32. Zhou, W., et al., Structure of the Human cGAS-DNA Complex Reveals Enhanced Control of Immune Surveillance. Cell, 2018. 174(2): p. 300–311 e11.

33. Dobbs, N., et al., STING Activation by Translocation from the ER Is Associated with Infection and Autoinflammatory Disease. Cell Host Microbe, 2015. 18(2): p. 157–68.

34. Zhang, C., et al., Structural basis of STING binding with and phosphorylation by TBK1. Nature, 2019. 567(7748): p. 394–398.

35. Zhao, B., et al., A conserved PLPLRT/SD motif of STING mediates the recruitment and activation of TBK1. Nature, 2019. 569(7758): p. 718–722.

36. Bjorgen, H. and E.O. Koppang, Anatomy of teleost fish immune structures and organs. Immunogenetics, 2021. 73(1): p. 53–63.

37. Schoetz, U., et al., Early senescence and production of senescence-associated cytokines are major determinants of radioresistance in head-and-neck squamous cell carcinoma. Cell Death Dis, 2021. 12(12): p. 1162.

38. Turnquist, C., et al., Radiation-induced astrocyte senescence is rescued by Delta133p53. Neuro Oncol, 2019. 21(4): p. 474–485.

39. Collin, G., et al., Transcriptional repression of DNA repair genes is a hallmark and a cause of cellular senescence. Cell Death Dis, 2018. 9(3): p. 259.

40. Saul, D., et al., A new gene set identifies senescent cells and predicts senescence-associated pathways across tissues. Nat Commun, 2022. 13(1): p. 4827.

41. Idda, M.L., et al., Survey of senescent cell markers with age in human tissues. Aging (Albany NY), 2020. 12(5): p. 4052–4066.

42. Widjaja, A.A., et al., Inhibition of IL-11 signalling extends mammalian healthspan and lifespan. Nature, 2024. 632(8023): p. 157–165.

43. Moiseeva, O., et al., Metformin inhibits the senescence-associated secretory phenotype by interfering with IKK/NF-kappaB activation. Aging Cell, 2013. 12(3): p. 489–98.

44. Moiseeva, V., et al., Author Correction: Senescence atlas reveals an aged-like inflamed niche that blunts muscle regeneration. Nature, 2023. 614(7949): p. E45.

45. Benayoun, B.A., et al., Remodeling of epigenome and transcriptome landscapes with aging in mice reveals widespread induction of inflammatory responses. Genome Res, 2019. 29(4): p. 697–709.

46. Salminen, A., et al., Activation of innate immunity system during aging: NF-kB signaling is the molecular culprit of inflamm-aging. Ageing Res Rev, 2008. 7(2): p. 83–105.

47. Li, X.D., et al., Pivotal roles of cGAS-cGAMP signaling in antiviral defense and immune adjuvant effects. Science, 2013. 341(6152): p. 1390–4.

48. Matthews, J.L., Common diseases of laboratory zebrafish. Methods Cell Biol, 2004. 77: p. 617–43.

49. Hopkins, J.W., et al., STING promotes homeostatic maintenance of tissues and confers longevity with aging. bioRxiv, 2024.

50. Martinez, J.C., et al., cGAS deficient mice display premature aging associated with loss of chromatin organization, derepression of LINE1 elements and induction of inflammation. Nat Aging, CO-SUBMISSION.

51. Zhen, Z., et al., Nuclear cGAS restricts L1 retrotransposition by promoting TRIM41-mediated ORF2p ubiquitination and degradation. Nat Commun, 2023. 14(1): p. 8217.

52. Liu, H., et al., Nuclear cGAS suppresses DNA repair and promotes tumorigenesis. Nature, 2018. 563(7729): p. 131–136.

53. Jiang, H., et al., Chromatin-bound cGAS is an inhibitor of DNA repair and hence accelerates genome destabilization and cell death. EMBO J, 2019. 38(21): p. e102718.

54. Wang, J., et al., The oxidative DNA lesions 8,5’-cyclopurines accumulate with aging in a tissue-specific manner. Aging Cell, 2012. 11(4): p. 714–6.

55. Yousefzadeh, M.J., et al., Tissue specificity of senescent cell accumulation during physiologic and accelerated aging of mice. Aging Cell, 2020. 19(3): p. e13094.

56. Tabula Muris, C., A single-cell transcriptomic atlas characterizes ageing tissues in the mouse. Nature, 2020. 583(7817): p. 590–595.

57. Gorgoulis, V., et al., Cellular Senescence: Defining a Path Forward. Cell, 2019. 179(4): p. 813–827.

58. Huang, W., et al., Cellular senescence: the good, the bad and the unknown. Nat Rev Nephrol, 2022. 18(10): p. 611–627.

59. Bhatelia, K., et al., MITA modulated autophagy flux promotes cell death in breast cancer cells. Cell Signal, 2017. 35: p. 73–83.

60. Bai, J. and F. Liu, cGAS‒STING signaling and function in metabolism and kidney diseases. J Mol Cell Biol, 2021. 13(10): p. 728–738.

61. Wischnewski, M. and A. Ablasser, Interplay of cGAS with chromatin. Trends Biochem Sci, 2021. 46(10): p. 822–831.

62. Li, Q., et al., Therapeutic Development by Targeting the cGAS-STING Pathway in Autoimmune Disease and Cancer. Front Pharmacol, 2021. 12: p. 779425.

63. Harel, I., D.R. Valenzano, and A. Brunet, Efficient genome engineering approaches for the short-lived African turquoise killifish. Nat Protoc, 2016. 11(10): p. 2010–2028.

64. Astre, G., et al., Genetic perturbation of AMP biosynthesis extends lifespan and restores metabolic health in a naturally short-lived vertebrate. Dev Cell, 2023. 58(15): p. 1350–1364 e10.

65. Kowarz, E., D. Loscher, and R. Marschalek, Optimized Sleeping Beauty transposons rapidly generate stable transgenic cell lines. Biotechnol J, 2015. 10(4): p. 647–53.

66. Bray, N.L., et al., Erratum: Near-optimal probabilistic RNA-seq quantification. Nat Biotechnol, 2016. 34(8): p. 888.

67. Wang, L., S. Wang, and W. Li, RSeQC: quality control of RNA-seq experiments. Bioinformatics, 2012. 28(16): p. 2184–5.

68. Love, M.I., W. Huber, and S. Anders, Moderated estimation of fold change and dispersion for RNA-seq data with DESeq2. Genome Biol, 2014. 15(12): p. 550.

69. Subramanian, A., et al., Gene set enrichment analysis: a knowledge-based approach for interpreting genome-wide expression profiles. Proc Natl Acad Sci U S A, 2005. 102(43): p. 15545–50.

70. Mootha, V.K., et al., PGC-1alpha-responsive genes involved in oxidative phosphorylation are coordinately downregulated in human diabetes. Nat Genet, 2003. 34(3): p. 267–73.

